# Crowding governs resistance confinement and guides the improvement of adaptive therapy in a 2D melanoma model system

**DOI:** 10.64898/2026.10.06.757145

**Authors:** Maximilian Eiche, Elias Ferdinand Fischer, Maja Milas, Timon Citak, Serhii Aif, Ana-Sunčana Smith, Jona Kayser

## Abstract

The emergence of treatment resistant mutants is one of the primary contributors to therapy failure in cancer. Evolution-based adaptive therapies aim to exploit the competition between sensitive and resistant cells to prolong therapeutic control. However, in the densely packed tissues of solid tumors, competition is not only determined by cell-intrinsic factors but also a result of local physical interactions between neighboring cells. The successful development of physics-informed adaptive therapies therefore hinges upon the bottom-up integration of tissue mechanics and experimentally grounded models of tumor evolution. Here, we show how population-level treatment response emerges from mechanical interactions between individual cells. Combining genetically tailored therapy-mimicry experiments in spatially structured 2D melanoma colonies with mechanistic, particle-based tissue simulations as digital twins in a real-to-sim-to-real framework, we reveal how treatment-induced changes in mechanical crowding shape a cascade of resistance expansion, spatial competitive escape, and consequent therapy failure. We then link these underlying physical mechanisms to population-level observables in an abstracted mathematical model for reinforcement-learning-based therapy optimization and a distilled treatment-decision-heuristic, which we test experimentally. Together, our results identify mechanical crowding and escape as governing principles of resistance-mediated therapy failure in solid tumours and provide a basis for mechanistically informed, evolution-based treatment strategies.

## Main

Cancer therapy often fails because heterogeneous tumour populations adapt under treatment, allowing resistant clones to expand [1, 2]. Adaptive therapy seeks to delay such failure by modulating selection rather than maximizing tumour reduction and has shown promise in preclinical and early clinical studies [3–5]. Its efficacy relies on ecological interactions between sensitive and resistant populations that are typically inferred from proxies of global tumour burden [6–8]. In solid tumours, however, spatial constraints and microenvironmental heterogeneity are likely to shape these interactions and influence clonal expansion [9, 10]. Understanding how local ecological processes, including emergent physical phenomena, govern system-level therapy dynamics and how clinically accessible observables can guide treatment decisions will be critical for the development of mechanistically grounded therapy strategies [6, 11].

The spatial organization of tumours is a key determinant of how competitive interactions unfold. In densely packed tissues, limited access to space and resources restricts interactions to local neighbourhoods, causing the expansion of resistant clones to depend not only on intrinsic fitness differences but also on spatial context [12–16]. Importantly, therapeutic interventions can actively reshape this context by changing growth pattern and resource availability, whereas withdrawal allows regrowth and re-establishment of competition [17]. These spatially structured and time-dependent effects introduce a dynamic coupling between treatment and clonal competition that is not captured by well-mixed descriptions, making their system-level consequences difficult to infer.

Despite growing recognition of the importance of local ecological interactions, a quantitative understanding of how they shape system-level treatment response remains limited [6]. Many experimental studies of adaptive therapy rely on bulk measurements of tumour burden, obscuring the spatial heterogeneity and local interactions that govern clonal dynamics [18, 19]. Although co-culture, spheroid, and organoid systems have begun to probe competitive suppression and aspects of tumour architecture, they provide limited opportunities for longitudinal, spatially resolved analysis of clonal competition across varying treatment schedules [20–22].

Spatially explicit mechanistic models provide one way to address this gap by linking local interaction rules, spatial constraints, mechanical feedbacks and proliferative dynamics to emergent population-level behaviour. Such models have shown that local interactions can qualitatively alter evolutionary outcomes and treatment efficacy [9, 17, 23–27]. However, their predictions remain difficult to validate experimentally.

At the same time, treatment optimization typically relies on reduced, low-dimensional state descriptions derived from aggregate measurements, because explicit spatial information is rarely available in clinical settings and may be impractical for strategy discovery [28, 29]. This creates a gap between the local scale at which competitive interactions occur and the global observables commonly used for treatment control. It remains unclear when the control-relevant consequences of spatial competition require explicit spatial resolution, and when they can be captured by simplified descriptions of system-level dynamics. Adequate abstractions are not only key for an accurate mechanistic understanding but are also the basis for the rapidly developing field of reinforcement-learning-based therapy optimization [29–31].

Here, we introduce a spatially structured *in vitro* melanoma system that enables long-term, single-cell-resolved tracking of resistant populations under temporally modulated therapy. By combining experiments and mechanistic spatial modeling, we investigate how local interactions shape resistance dynamics. Distilling the uncovered mechanisms into a context-constrained effective model together with reinforcement learning, we show how crowding-mediated competition can inform physics-guided evolution-based therapy strategies.

## Results

To investigate how global therapy dynamics and local competition mechanisms are connected across scales (Fig. 1a), we developed an integrated *in vitro*/*in silico* framework that combines long-term tracking of whole-colony treatment-sensitive and resistant cell populations under time-varying treatment landscapes in experiments with interpretable mechanistic modeling of the mechanical interactions between individual cells.

**Figure 1:**
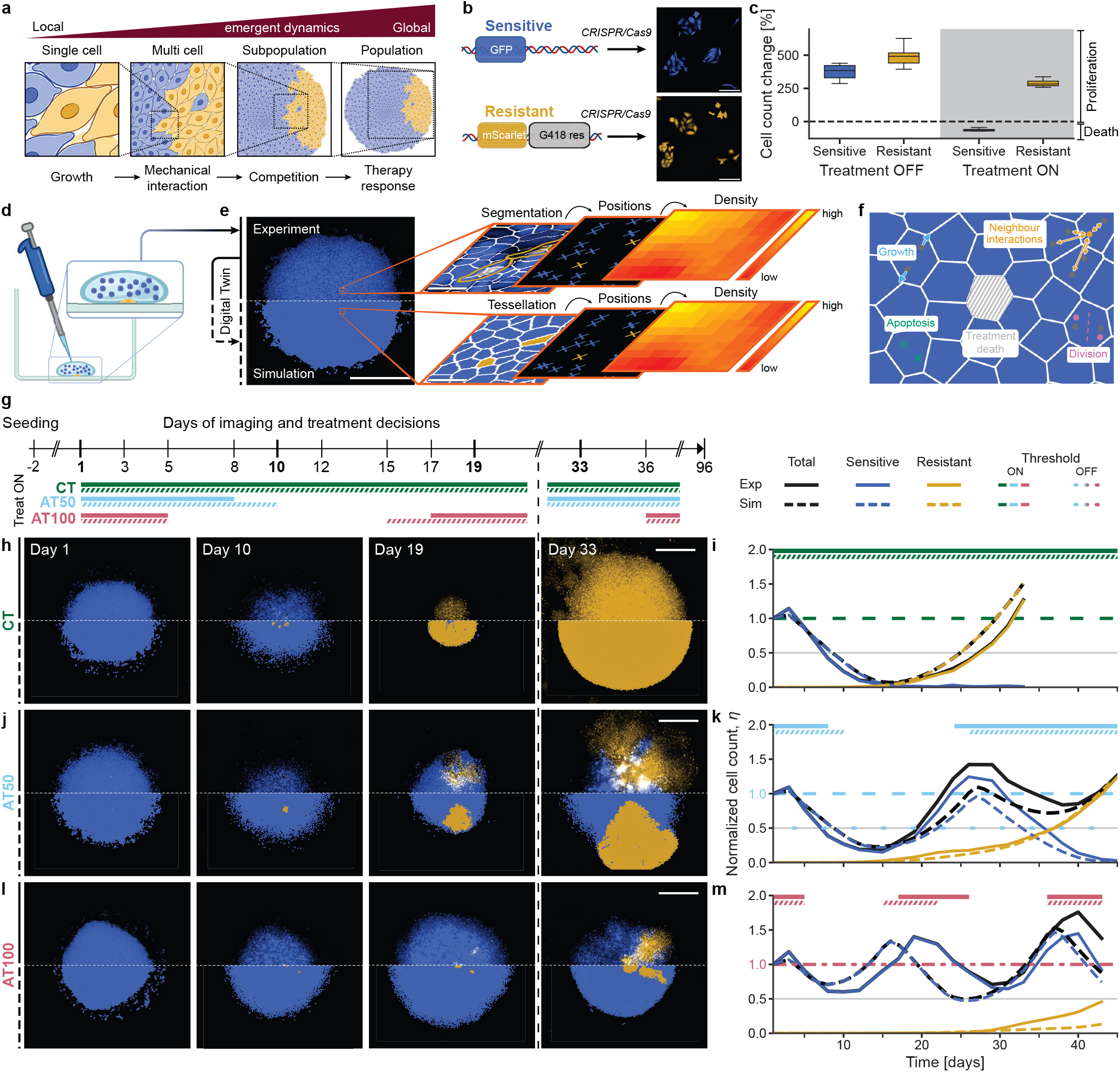
An integrated experiment-simulation system to study the physics of adaptive therapy. **a** Schematic of how system-level phenomena emerge from local interactions. **b** Genotype schematic and example images of synthetically engineered G418-resistant (yellow) and sensitive (blue) melanoma cells. Scale bar, 100 μm. **c** Type-specific treatment effects as measured by change in cell count after 3 days. Boxes show median and interquartile range (IQR); whiskers indicate 1.5× IQR, n = 8 replicate wells per cell line and condition. **d** Illustration of colony seeding. **e** Example of whole-colony, single-cell resolution fluorescence image (scale bar 2 mm, gamma = 0.7) with schematic segmentation, position-map and generation of density maps *in vitro* (scale bar, 50 μm). Bottom row: Dissipative particle dynamics simulations serve as digital twins that are initiated from experimental-derived cell positions, following the same analysis pipeline. **f** Schematic of the particle-based simulation framework. **g** Timeline indicating imaging, medium exchange and treatment decision days. Bold timepoints correspond to images below. Bars show periods of treatment for CT (green), AT50 (blue) and AT100 (red) replicates *in vitro* (solid) and *in silico* (striped). **h, j, l** Example images of colonies under CT (h), AT50 (j) and AT100 (l) *in vitro* and matched *in silico*. Scale bar, 2 mm, blue channel: gamma = 0.1, yellow channel: gamma = 0.2. Blue and yellow indicate sensitive and resistant cells, respectively, while white denotes overlap. **i, k, m** Corresponding total (black), sensitive (blue) and resistant (yellow) cell count *η* (normalized to initial total) of CT (i), AT50 (k) and AT100 (m) colonies *in vitro* (solid) and *in silico* (dashed). Horizontal lines give the treatment thresholds of each strategy; bars above give treatment periods. Parts of panel b and d illustrations were created in BioRender [32].

The experimental system is based on two CRISPR-engineered, fluorescently barcoded human melanoma cell lines of which one is G418-sensitive (EGFP, pseudo-coloured in blue) while the other is resistant (mScarlet, pseudocoloured in yellow)(Fig. 1b, c). To mimic one *de novo* mutation event, we seeded individual resistant cells surrounded by a large sensitive population, forming densely-packed colonies with a defined boundary (Fig. 1d). We subsequently conducted highly-parallelized longitudinal therapy-mimicry experiments for up to 96 days with single-cell resolution, whole-colony images acquired three times per week.

For closed-loop adaptive treatment decisions, same-day image analysis was performed via a custom automated analysis pipeline (Fig. 1e). In short, fluorescence images were segmented into individual cells, extracting the type and center of mass position for each cell [33–35]. Based on this data, we computed type-specific population sizes and spatial density maps.

To resolve and understand the mechanobiological constraints governing resistant cell expansion, we complemented experiments with dissipative particle dynamics simulations [23]. Acting as digital twins, these simulations have been shown to quantitatively capture tissue development [23, 27]. In this mechanically explicit framework, each cell is represented by two particles and incorporates cell growth, interactions with neighbouring cells [23], a division rate regulated by cell size [27, 36], lifetime-dependent stochastic apoptosis, and therapy-induced death determined by cell sensitivity and drug absorption capacity (Fig. 1f, Supplementary Methods Section 4.1). Importantly, key population-level phenomena, such as competition and therapy response, emerge intrinsically from the underlying local dynamics. Following global parameterization, each digital twin was initialized from an experiment-derived colony configuration and subjected to the same treatment strategies, based on simulated cell counts (Fig. 1e, f, Supplementary Methods Section 4.2).

Using this platform, we conducted matched *in vitro*/*in silico* experiments to compare continuous therapy (CT) and adaptive therapies (AT) (Fig. 1g, Supplementary Methods Section 4.3). For continuous therapy, a constant drug dose was maintained over the entire course of the experiment. For adaptive strategies, treatment was paused when the total cell count dropped either below 50% (AT50) or 100% (AT100) of the initial cell count and resumed upon regrowth above baseline.

In both *in vitro* and *in silico* the CT baseline strategy yields complete eradication of the sensitive population, followed by rapid growth of resistant cells (Fig. 1h, i). For AT50, we find a delayed rebound of sensitive cells that, while slowing resistance growth, is typically insufficient to maintain resistance confinement for more than one treatment cycle (Fig. 1j, k). Only AT100 exhibits a more pronounced effect on resistance suppression, often maintaining control over several treatment cycles (Fig. 1l, m).

Comparing population-level dynamics by longitudinal measurements of type-specific cell numbers we find that the independently calibrated simulation captures the experimental observations both qualitatively (Fig. 1h, j, l) and quantitatively (Fig. 1i, k, m).

However, the suppressing effects of sensitive cell regrowth during adaptive strategies remain limited, especially for AT50, warranting a closer investigation of how spatial population structure affects therapy response.

### Spatial escape of resistant clones drives loss of therapeutic control

One inherent consequence of a spatially bounded population is that resistant cells reaching the colony edge are no longer subject to local competition with sensitive cells even if the total count rebounds. We will refer to this form of spatial competitive release as “escape” (Fig. 2a). Delaying escape might be the key objective to facilitate long-term resistance control.

**Figure 2:**
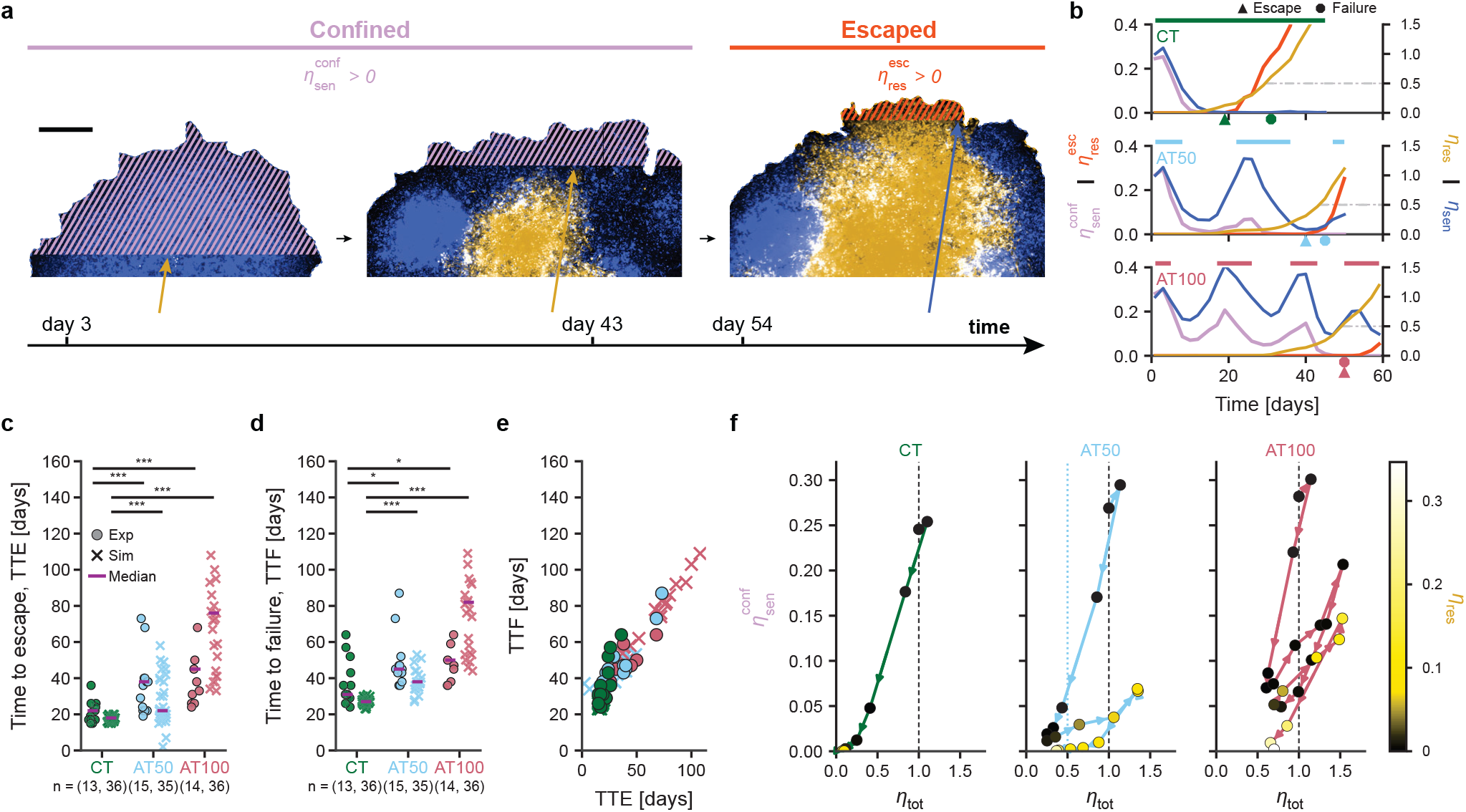
Control of resistant clones depends on clonal escape dynamics. **a** Representative fluorescence images illustrating escape of a resistant clone over time. Striped areas indicate the region of confining sensitive cells (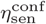, purple), and escaped resistant cells 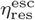 orange). Arrows indicate the most forward resistant or sensitive cell. Scale bar = 200 μm, gamma = 0.3. **b** Quantification of representative escape dynamics over time for CT, AT50, and AT100. Horizontal lines above indicate treatment. Gray lines indicate treatment failure threshold. **c** Time-to-escape 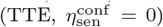 for experiments and simulations. Points are individual colonies, bars indicate median. Numbers below groups indicate respective *n*. Statistical tests: two-sided log-rank tests. Non-event and truncated observations are not shown and treated as right-censored. **d** Time-to-failure (TTF, *η*_res_ *≥* 0.5), analogous to c. **e** Correlation between TTE and TTF (*in vitro*: Pearson’s *r* = 0.85, *p* = 5.1 10^*−*10^; *in silico*: *r* = 0.98, *p* = 4.1 *×* 10^*−*49^). **f** Visualization of ratchet-like clonal escape trajectories. Marker colors indicate the corresponding normalized resistant cell burden, *η*_res_. Dashed lines indicate initial values, and the dotted line denotes the AT50 treatment-off threshold.

To study the dynamics of resistance escape more systematically, we quantify the degrees of confinement by the number of sensitive cells between the resistant clone and open space before escape,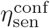, and the degree of escape by the number of resistant cells more forward than any sensitive cells, 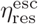(Fig. 2a, b).

The dynamics of the confining sensitive layer reveal a key difference between both adaptive strategies. For AT100, 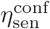 deteriorates during the initial round of treatment but rebounds before the second cycle. In contrast, AT50 shows only marginal rebound, despite both strategies recovering to the initial total population size. This decrease of confinement layer rebound can also be observed for AT100, but only at a later stage *in vitro* as well as *in silico* (Supplementary Fig. 6).

To probe how confinement dynamics relate to treatment success, we measured the time to escape 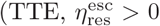; Fig. 2c) and the time to failure of resistance control (TTF, *η*_res_ *≥* 0.5; Fig. 2d), for each strategy in both *in vitro* and *in silico*. Compared with CT, AT100 significantly extended both TTE and TTF in experiments and simulations (Exp: ΔTTE = 23 days; 95% CI = 7–49 days; *p* = 1.13 *×* 10^*−*5^; ΔTTF = 19 days; 95% CI = 2–33 days; *p* = 0.0138; Sim: ΔTTE = 58 days; 95% CI = 42–75 days; *p* = 5.29 *×* 10^*−*19^; ΔTTF = 55 days; 95% CI = 47–67 days; *p* = 2.31*×* 10^*−*16^). AT50 likewise significantly extended TTE and TTF relative to CT in both experiments and simulations, although the simulated delay in TTE was smaller than that observed for AT100 (Exp: ΔTTE = 16 days; 95% CI = 0–23 days; *p* = 4.02*×*10^*−*4^; ΔTTF = 14 days; 95% CI = 0–21 days; *p* = 0.0357; Sim: ΔTTE = 4 days; 95% CI = 2–14 days; *p* = 9.07*×* 10^*−*8^; ΔTTF = 11 days; 95% CI = 8–17 days; *p* = 1.75*×*10^*−*13^).

For all strategies, TTE was strongly correlated with TTF (Pearson’s *r* = 0.847, 95% CI = 0.667–0.930, *p* = 5.1 *×* 10^*−*10^, *n* = 33 *in vitro*; *r* = 0.979, 95% CI = 0.954– 0.991, *p* = 4.1 *×* 10^*−*49^, *n* = 71 *in silico*), corroborating our hypothesis without establishing causality (Fig. 2e).

Supported by high-resolution *in silico* data (Supplementary Fig. 5), these findings suggest that the observed dynamics of the confining sensitive cell layer result from a two-sided spatial ratchet: Treatment deteriorates confinement by both peripheral killing of sensitive cells while also triggering resistance expansion (Fig. 2f). Both processes together cause a decrease of 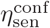.However, while the decrease in sensitive cell number is reversible, resistance expansion is not. Since sensitive rebound is capped by the total burden threshold, this irreversible ratchet will gradually cause confinement deterioration and eventually escape.

### Resistant growth suppression is governed by local crowding

Going beyond the limits accessible experimentally, our simulations allow us to directly probe how the observed population dynamics emerge from interactions at the level of individual cells, where proliferation is modulated by the mechanical constraints exerted by neighboring cells (Fig. 3a). During the expansion phase of the cell cycle, each cell has to extend against these external forces, decreasing its expansion rate (*λ*) and delaying the entry into the division phase (Fig. 3b) [27]. Relating *λ* to the local total cell density *ρ*_tot_, a measure also available in experiments, shows that higher densities are strongly associated with decreased cell expansion (Fig. 3c, Supplementary Methods Section 4.4). This is consistent with previous work on mechanosensitivity of cell proliferation [23, 27, 36, 37]. We can therefore use density as a proxy for crowding-mediated mechanical constraints on cell growth.

It should be noted that the effects of density can also be cell type dependent, and that in our parameterization the relation between *λ* and *ρ*_tot_ for sensitive cells differs from that of resistant cells (Supplementary Table S1). However, we will here focus on density effects on resistance growth.

The mechanical effects on cell growth are clearly reflected in spatial expansion and contraction of resistant and sensitive subpopulations, which we quantify by computing spatially-resolved density maps *ρ*_res_(*x, y*) and *ρ*_sens_(*x, y*) alongside associated density-change maps Δ*ρ*_res_(*x, y*) for both simulations and experiments (Fig. 3d, e). Together, these reveal anti-correlated dynamics in which resistance expansion is arrested in high-density environments but can proceed when crowding is reduced due to treatment-induced depletion of sensitive cells (Fig. 3f, g). Despite substantial spatial and temporal heterogeneity across different treatment patterns, this relationship can be quantitatively summarized by a master curve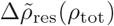 that collapses the spatial maps into a crowding-growth axis (Fig. 3h, i). For all strategies, high-density regions were associated with resistant stasis, whereas low-density regions permitted growth.

**Figure 3:**
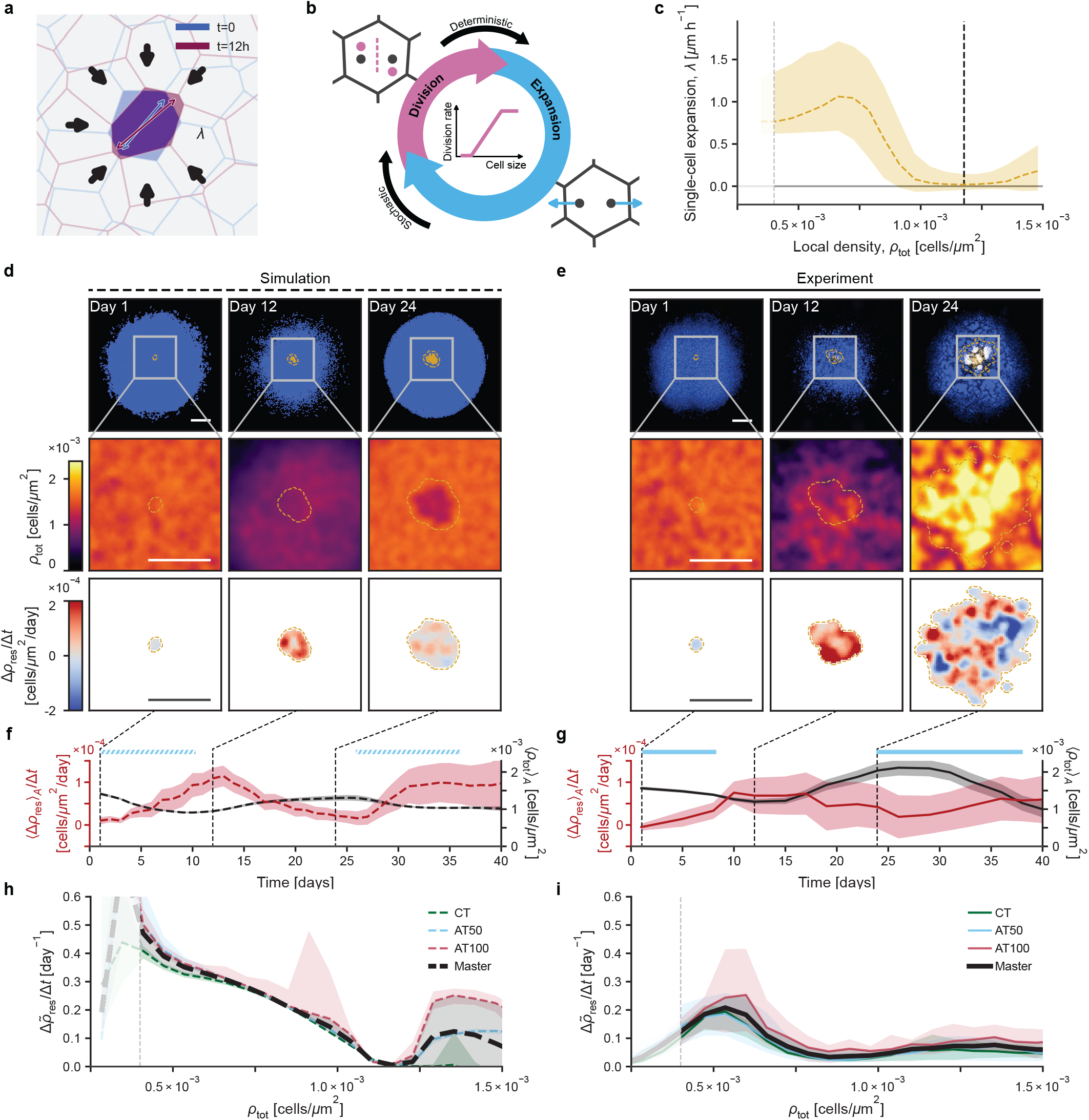
Local crowding governs the suppression of resistant growth. **a** Schematic of mechanical forces countering cell expansion in the simulation. **b** Schematic illustrating the simulated cell cycle. **c** Single cell expansion *λ* as a function of total local density *ρ*_tot_ for resistant cells *in silico*. Dashed yellow curve shows median, the shaded region the interquartile range. *ρ* values below 0.4 *×* 10^*−*3^cells*/μ*m^2^ correspond to non-confluent states and are excluded. Dashed black vertical line indicates homeostatic pressure. **d, e** Representative *in silico* (d) and *in vitro* (e) colonies, coarse-grained total density *ρ*_tot_ and resistant density-change Δ*ρ*_res_/Δ*t* maps for low and high density time points. Yellow dashed outlines indicate pixels with sufficient local resistant-cell density (threshold = 0.05 cells*/*1,000 *μ*m^2^). Blue channel: gamma = 0.45, yellow channel: gamma = 0.2. **f, g** Corresponding area-averaged resistant density change (Δ*ρ*_res_) _*A*_*/*Δ*t* (red, left axis) and total density *(ρ*_tot_) _*A*_ (black, right axis), running mean with SD (shaded). Bars above give treatment periods, dashed lines mark the days shown in d and e. **h, i** Binned relative resistant density change 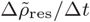 as a function of *ρ*_tot_ *in silico* (h) and *in vitro* (i) for CT, AT50 and AT100, per-strategy median with interquartile range (shaded), and the mean over strategies (black) with SD (shaded). *ρ* values below 0.4 *×* 10^*−*3^cells*/μ*m^2^ correspond to non-confluent states.

Together, these observations suggest that density might indeed be used as an effective control parameter for therapeutic decisions.

### Local crowding-mediated competition can be abstracted into a population-level model for RL-guided therapy optimization

Because spatially resolved measurements of crowding are rarely available for treatment decisions in a clinical context, we asked whether the outcome relevant dynamics could be further abstracted into a coarse-grained model based on more accessible population-level observables without loosing predictive power on strategy optimality.

Finding that normalized total-cell number *η*_tot_ closely tracked total-cell density*(ρ*_tot_)_*A*_ in experiments before escape (Fig. 4a), using experimental data (Fig. 4b), we constructed a minimal non-spatial effective model based on an established two-population generalized Lotka–Volterra (GLV) system (Fig. 4c). The calibrated GLV model reproduced the qualitative pre-escape population dynamics for AT100. Importantly, AT100 consistently outperformed AT50 and CT across the GLV model, the spatial simulations and the experiments (Supplementary Fig. 7). Thus, while the GLV model should not be interpreted as a faithful quantitative description of the complete spatial dynamics, it represents a control-relevant abstraction of the pre-escape regime in which competition can still be exploited. As a direct consequence, the GLV model may be used as a computationally efficient environment surrogate for treatment optimization beyond fixed-threshold strategies.

**Figure 4:**
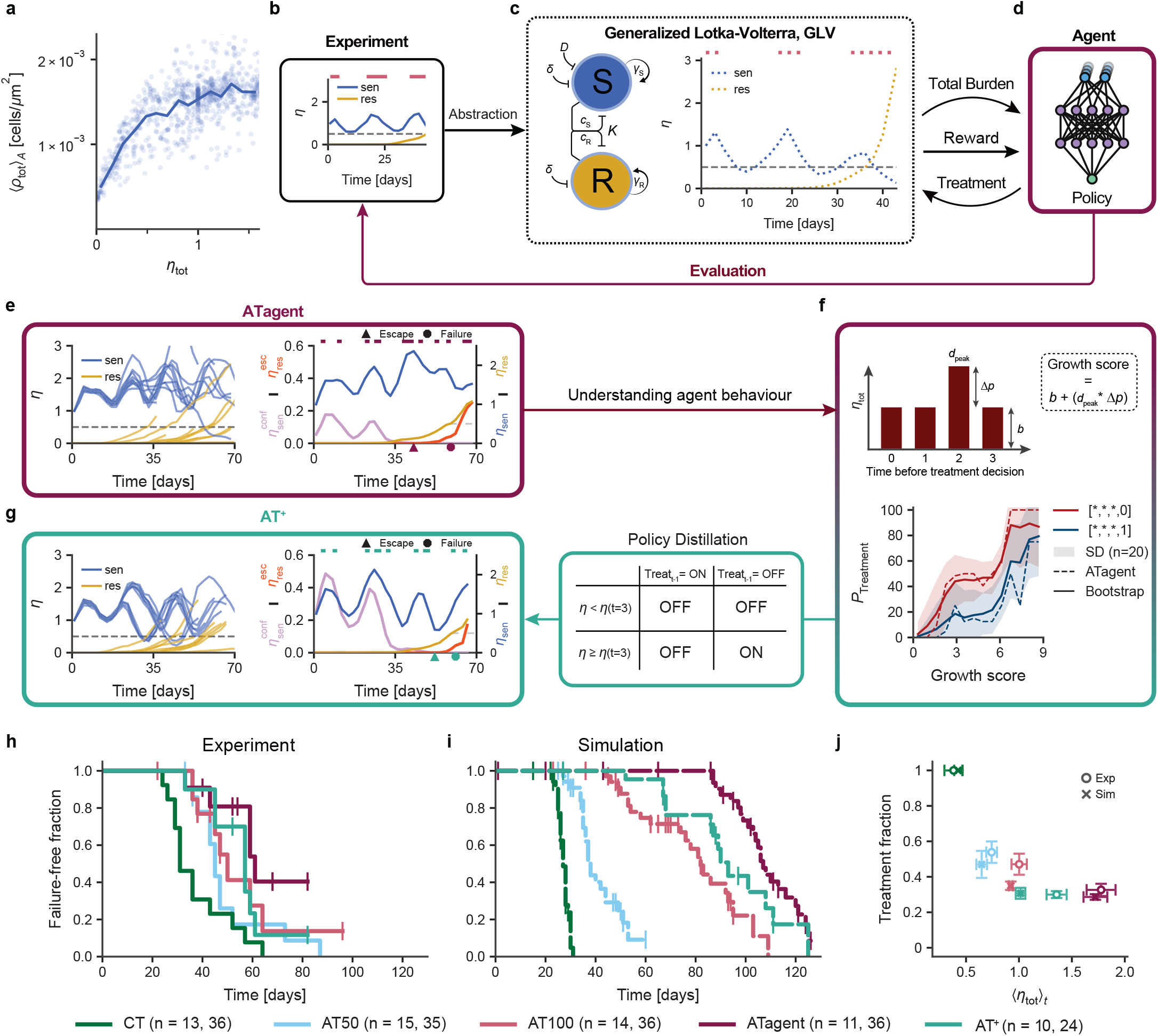
Optimized evolutionary treatment strategies prolong survival *in vitro* and *in silico*. **a** Mean normalized total-cell number *η*_tot_ and area-averaged total-cell density in resistant clone region *(ρ*_tot_ )_*A*_ before escape. **b** Example *in vitro* normalized cell count with dashed horizontal bars above indicating treatment. Horizontal gray line indicating treatment failure threshold. **c** Effective two-population GLV model of sensitive (S) and resistant (R) cells, with a representative GLV fit to the AT100 example in b. Dashed bars above indicate treatment periods. Horizontal gray line indicating treatment failure threshold. **d** Schematic of agent trained on the GLV environment in b. **e** *In vitro* normalized cell count trajectories under ATagent, with a representative example showing confining sensitive 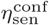 and escaped resistant cells 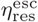. Horizontal gray line indicating treatment failure threshold. **f** Retrospective policy-behaviour analysis of 20 PPO agents trained on bootstrap-refitted effective models. Shading indicates *±* 1 sample SD. Synthetic four-observation burden histories are summarized by the growth score *G* = *b* + (*d*_*peak*_ *∗* Δ*p*), with baseline burden *b*, peak magnitude Δ*p*, and peak position *d*_*peak*_ = 0,…, 3 from oldest to most recent. **g** Qualitative and quantitative depiction of the colony dynamics under the distilled AT^+^ strategy as in e. **h, i** Failure-free fraction per strategy *in vitro* (h) and *in silico* (i); ticks mark censored replicates, n = (*in vitro, in silico*). **j** Treatment fraction against time-averaged normalized total burden *η*_tot *t*_ per strategy *in vitro* and *in silico*. Bars indicate SD. n = (*in vitro, in silico*).

More specifically, this abstracted model can serve as the training environment for reinforcement learning (RL), which optimizes observation-based treatment decisions to maximize a predefined reward function, such as long-term outcome (Fig. 4d). The advantage of such an unbiased approach allows control principles to naturally emerge from the system dynamics rather than being confined to pre-conceived notions.

The RL agent was trained in a fixed effective-model environment using proximal policy optimization and two sequential reward functions. At each decision point, the agent observed only the normalized total population burden *η*_tot_ at the four most recent measurements together with the corresponding treatment decisions and selected whether treatment should be applied during the next interval. It had no access to clonal composition, local density or spatial position. To assess the robustness of the learned decision logic to uncertainty in model calibration, we subsequently refitted the effective model to 20 stratified bootstrap datasets and trained one additional policy for each refit. The policy selected for experimental scheduling and validation is hereafter termed ATagent (Fig. 4e).

To identify which features of the observation history governed treatment decisions, we evaluated the trained policies across a predefined set of synthetic inputs (Fig. 4f, Supplementary Methods Section 5.3). The fraction of deterministic treatment-on decisions could not be reduced to current burden alone. Across the input ensemble, treatment was selected more frequently when burden had risen over the preceding observations rather than depending only on its instantaneous value, and more frequently after a treatment-free interval than when therapy had already been applied recently. We distilled this behaviour into a simplified decision heuristic, AT^+^, which applies treatment only if it was withheld at the preceding decision point and burden exceeds treatment threshold. For consistency with treatment initiation of ATagent, we defined the treatment threshold as the total burden on day 3.

To test whether this treatment logic transferred beyond the effective model used for training, we applied ATagent and AT^+^ to *in vitro* colonies and corresponding digital twins, recording population-level metrics as before. Both strategies adhered to the experimental measurement time points and used total burden as their only population-level input. Representative trajectories reveal the same qualitative pulse-pause dynamics in experiments and spatial simulations (Fig. 4g). ATagent and AT^+^ delivered short pulses separated by treatment-free intervals during which sensitive cells and total burden could recover. Importantly, corresponding 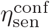 trajectories remained more stable than under AT50 and AT100, delaying the eventual confinement collapse (Supplementary Fig. 6). Although ATagent was never provided with density measurements, its pulse pattern - and consequently those of AT^+^ - avoided prolonged low-burden states associated with reduced local density. Finally, comparison of all strategies at the population level confirmed this advantage in both experimental colonies and their digital twins (Fig. 4h and i). Kaplan–Meier curves showed delayed failure from CT through AT50 and AT100 to the model-derived strategies. Median time to failure increased from 31 days (CT) to 45/50 days (AT50/AT100) and 57/61 days (AT^+^/ATagent) *in vitro*, with an even stronger separation *in silico* (27, 38, 82, 93, and 107 days, respectively). CT versus ATagent differed in both settings (Exp, *p* = 0.0052; Sim, *p* = 5.9 *×* 10^*−*15^; two-sided log-rank tests with Bonferroni correction). Extending the evaluation beyond disease control alone, we compared all strategies in an adversity space defined by the percentage of days under treatment and the normalized total burden *η*_tot_ (Fig. 4j). In this joint comparison, AT^+^ achieved a more favorable balance than ATagent, with similar treatment exposure and a lower time-averaged normalized population burden (median burden difference, AT^+^ minus ATagent: Exp, *−*0.44, 95% CI [*−*0.51, *−*0.31], *p* = 0.00012; Sim, *−*0.74, 95% CI [*−*0.77, *−*0.62], *p* = 9.6 *×* 10^*−*11^).

## Discussion

Evolution-based cancer therapy seeks to control resistance by maintaining competition between drug-sensitive and resistant cells. Our study connects this ecological strategy to tissue biophysics by showing how competition depends on the local physical environment that cells create and treatment continually reshapes. By combining longterm, spatially resolved experiments in melanoma colonies with mechanically explicit digital twins, we uncover how population-level treatment response emerges as an inherent consequence of local mechano-competition. Closing the real-to-sim-to-real loop, we then abstract these physics-driven dynamics into an effective model for RL-guided therapy optimization and experimentally demonstrate improved treatment outcomes over previous AT strategies.

Three new perspectives emerge from this work. First, understanding resistance evolution as an emergent phenomenon in proliferating active matter systems, yields crucial insights on how physical mechanisms shape therapy response. Mechanical feedback has long been proposed to coordinate tissue growth, and applied compression can arrest proliferation in epithelial tissues and tumour spheroids [27, 38–41]. This work now demonstrates how such mechano-feedback becomes a treatment-modulated competition mechanism in solid tumors. This interpretation extends ecological accounts of competitive release [3, 7] by providing a physical counterpart to previous demonstrations that spatial structure can change mutant selection and adaptive-therapy performance [12– 14, 42].

Second, the central geometric consequence is an irreversible escape ratchet. Treatment both thins the confining sensitive-cell layer separating a resistant clone from open space and permits the clone to expand into the resulting low-density region. Even if the total cell number can recover during a treatment pause, the confining margin does not. Once the clone reaches the colony boundary, competition becomes effectively one-sided and global regrowth of sensitive cells no longer restores confinement. The close association between escape and failure therefore identifies boundary crossing as a macroscopic spatial loss of control, consistent with confinement-and-release dynamics driven by other treatment-modulated landscapes [15, 17].

Third, and maybe most intriguingly, detailed information of the underlying local state variables, such as mechanical constraints, need not always enter a treatment controller explicitly to be effective. The only requirement is a global observable, like established measures of total burden, that can serve as an abstracted proxi for the local state variable within the optimization-relevant pre-escape phase. In this sense, the RL-based optimization and distillation implicitly identified a physical control principle rather than merely a schedule. Importantly, rather than negating the critical relevance of physical mechanisms, these findings offer a mechanistic basis for burden-guided adaptive-therapy models [6, 28]. The integrated real-to-sim-to-real approach complements model-based optimization studies [30] by experimentally testing an interpretable policy in the spatial system from which its effective training environment was abstracted.

There are several limitations that need to be considered when interpreting our findings. For example, density was measured directly, but mechanical pressure was not. Agreement with the mechanically explicit simulations shows that pressure-modulated growth is consistent with, and sufficient to reproduce, the observations. However, it does not establish it as the unique micro-scopic cause. Biochemical contact inhibition, mechanosensitive signalling, cell shape, paracrine interactions or other density-correlated processes could contribute to the same growth law [9, 41, 43]. Moreover, resistance was genetically fixed, whereas maintaining a large sensitive population in a real tumour may also increase the supply of new resistant variants. The two-dimensional, sharply bounded colonies omit three-dimensional architecture, extracellular matrix, stromal and immune interactions, active invasion, and spatial gradients in nutrients and drug. Their single-lesion setup also cannot represent metastatic sites with distinct compositions and exposure. Testing the proposed control principle will therefore require direct force or mechanotransduction readouts, lineage-resolved mutation assays, and three-dimensional or multi-lesion systems in which crowding and boundary access can be perturbed independently. These extensions should establish how well the principles uncovered here generalize to increasingly complex pre-clinical settings and, eventually, patients.

More broadly, this work demonstrates the value of understanding how population-level therapy dynamics emerge from fundamental physical interactions across scales for the rational design of improved physics-informed adaptive therapy strategies.

## Methods

### Cell culture

The parental human melanoma cell line Mel JuSo was obtained from the laboratory of Prof. Anja Boßerhoff. Cells were maintained in RPMI 1640 medium (PAN-Biotech, P04-18047) supplemented with 10% fetal bovine serum at 37 ^*°*^C in a humidified atmosphere containing 8% CO_2_. Cells were passaged two to three times per week upon reaching approximately 60–80% confluence. Cultures were routinely tested for mycoplasma contamination.

### Plasmid construction

Two fluorescently labelled Mel JuSo derivatives with differential G418 sensitivity were generated by CRISPR/Cas9-mediated homology-directed integration at the AAVS1 safe-harbour locus. The AAVS1-targeting plasmid pME_001 was eSp-Cas9(1.1)_No_FLAG_AAVS1_T2 (Addgene plasmid #79888; RRID:Addgene_79888), which encodes eSpCas9(1.1) and the AAVS1-T2-targeting sgRNA 5’-GGGGCCACTAGGGACAGGAT-3’ (Supplementary Fig. 8). The donor plasmid used to generate the sensitive cells, pME_002, was the unmodified pAAVS1-P-CAG-GFP (Addgene plasmid #80491; RRID:Addgene_80491) (Supplementary Fig. 9). This plasmid contains AAVS1 homology arms, a CAG-driven EGFP cassette, and a promoterless splice-acceptor-T2A-puromycin-resistance cassette.

The donor plasmid used to generate the resistant cells, pME_016, was constructed by Gibson assembly using the empty AAVS1 donor backbone pAAVS1-P-MCS (Addgene plasmid #80488; RRID:Addgene_80488), an mScarlet-H-containing fragment amplified from pmScarlet-H_C1 (Addgene plasmid #85043; RRID:Addgene_85043), and a CMV-driven neomycin phosphotransferase cassette amplified from pTS2404-Tier1-PhCMV-NeoR (Addgene plasmid #169496; RRID:Addgene_169496) (Supplementary Fig. 10). Fragments were amplified using Q5 Hot Start High-Fidelity 2X Master Mix and the primers listed in Supplementary Table S2 and assembled using NEBuilder HiFi DNA Assembly Master Mix. Candidate colonies were screened by colony PCR, and the completed pME_016 construct was verified by Sanger sequencing. The resulting donor retained the AAVS1 homology arms and promoterless puromycin-selection cassette and additionally encoded mScarlet-H and NeoR, thereby conferring G418 resistance.

### Generation and validation of sensitive and resistant cell lines

Approximately 2 *×* 10^5^ Mel JuSo cells were seeded per well in six-well plates one day before transfection. Cells were transfected using Lipofectamine 3000 and P3000 reagent in Opti-MEM according to the manufacturer’s protocol. Per well, 0.75 *μ*g pME_001 and 1.75 *μ*g of either pME_002 or pME_016 were combined with 3.75 *μ*l Lipofectamine 3000 . Complexes were incubated for 10– 15 min at room temperature before being added dropwise to the cells. Cas9-only, donor-only, and no-DNA controls were included.

Puromycin selection at 2 *μ*g/ml was initiated 48 h after transfection. Surviving reporter-positive cells were gradually expanded and single-cell sorted by fluorescence-activated cell sorting to establish clonal populations. Because reduced mScarlet fluorescence was subsequently observed in the resistant cells, these cells underwent a second round of single-cell sorting. Their resistant phenotype was further consolidated through stepwise G418 dose escalation up to 4 mg/ml. Resistant cells were subsequently maintained in medium containing 0.5 mg/ml G418. Selection antibiotics were omitted after seeding mixed-population experiments, in which G418 was applied exclusively according to the assigned treatment schedule. Genomic DNA was extracted using QuickExtract DNA Extraction Solution (Merck, LGCQE09050). Site-specific integration was assessed by PCR using the primer pairs listed in Supplementary Table S2. Integration in the resistant cells was assessed using an insertion-control PCR with oME_068/oME_264 and a PCR spanning the NeoR sequence with oME_068/oME_175. Products of the expected sizes confirmed integration of the respective donor constructs at the AAVS1 locus. The three highest-ranked potential off-target loci predicted using CRISPOR were amplified using the primers listed in Supplementary Table S2 [44].

For cryopreservation, exponentially growing cells were resuspended at approximately 1*×* 10^6^ cells per vial in freezing medium containing 60% RPMI, 30% fetal bovine serum, and 10% DMSO. Vials were cooled overnight at *−* 80 ^*°*^C in a controlled-rate freezing container and subsequently transferred to liquid nitrogen.

### AlamarBlue viability and drug-response assay

Metabolic activity was quantified using AlamarBlue Cell Viability Reagent (Thermo Scientific, DAL1025). Calibration curves covering 0–6,400 cells per well were generated for each cell line using six technical replicates per density. Cell-free wells were included for background correction, and Rhodamine B wells were included to calibrate measurements between plates. Outer wells were excluded to reduce edge effects. AlamarBlue was added to a final concentration of 10%, followed by incubation for 2 h at 37 ^*°*^C and fluorescence measurement at 554 nm excitation and 593 nm emission.

For G418-response measurements, sensitive and resistant cells were seeded at 1,000 cells per well in 100 *μ*l medium. G418 concentrations of 0, 0.025, 0.05, 0.1, 0.2,, 0.4, 0.6, 0.8, 1, 2, and 5 mg/ml were tested using eight technical replicates per cell line and concentration. A pretreatment measurement was performed two days after seeding, after which G418 treatment was initiated. Further measurements were performed after 2, 5, and 7 days of treatment. At each time point, AlamarBlue was added directly to the existing culture medium, including the respective G418 concentration. Following measurements on treatment days 2 and 5, the medium was exchanged and G418 was replenished. Control experiments confirmed that G418 itself did not affect the AlamarBlue fluorescence readout.

Fluorescence values were background-corrected, normalized to the corresponding RhodamineB calibration signal, and subsequently normalized to the pretreatment measurement of the respective condition. Drug-free proliferation was assessed separately using 16 technical replicates per cell line, with measurements performed 2, 5, and 7 days after seeding. The resulting differential G418-response profiles are shown in Supplementary Fig. 2.

Cryopreserved cells were thawed and cultured for 3 passages prior to 2D radial growth experiments for cells to reach a healthy and sustainable growth. Washed and trypsinized, resistant cells were harvested, centrifuged for 3 min at 300g and then resuspended in cell culture medium at a density of 1333 cells/ml. Single drops of 3 μl resistant cell suspension were seeded in the center of a well of a 24-well plate. Cells were allowed to settle for 4-5 hours under normal cell culture conditions. Then washed and trypsinized, sensitive cells were harvested, centrifuged for 3 min at 300g and then resuspended in cell culture medium at a density of 1 10^6^ cells/ml. Drops of 20 μl sensitive cell suspension were added to the prior seeded resistant cells. Cells were allowed to settle for 18 hours under normal cell culture conditions. Fresh media was then added to cover the whole well. At this point the cells formed a round colony with a radius of 4-5 mm and 20,000 cells per colony (Supplementary Fig. 3). Consequently, the resistant cells were initially localized near the center of the colony (Supplementary Fig. 4).

### Image analysis and feedback control

Colonies were imaged on Mondays, Wednesdays, and Fridays using a Zeiss LSM 900 microscope controlled by ZEN software. Widefield fluorescence images were acquired using a 5 *×* objective (NA 0.16) with an Axiocam 705 camera and a 0.63*×* camera adapter. Green and red fluorescence were acquired using the EGFP and mScarlet channels, respectively. Images were acquired at a calibrated pixel size of 1.095 *μ*m*/*px. Multiple image tiles covering each colony were stitched using the Stitching tool in ZEN. Sensitive and resistant fluorescence channels were segmented independently using custom Cellpose models [33–35]. Each accepted object was represented by the centre of mass of its segmentation mask. To correct for detector saturation, high-intensity regions not assigned to segmented objects were identified and their missing cell number was estimated from the saturated area using a calibration obtained from non-saturated images. Corrected total counts were used for treatment decisions and downstream population analyses. Model versions, segmentation parameters and saturation-correction details are provided in Supplementary Methods Section 1.

For CT, G418 was maintained throughout the experiment. For adaptive treatment, the corrected total population size was normalized to the initial count of the individual colony. AT50 was applied from the start and paused once the total count fell below 50% of baseline; treatment resumed when the population returned to baseline. AT100 was paused once the total count fell below baseline and resumed on return to baseline. G418 was applied at 0.4 mg ml^*−*1^. Experiments were ended for an individual colony when its total population reached 300% of baseline.

### Spatial escape and density analyses

Resistant clones and the sensitive population were followed across successive images. Confinement and escape were quantified by comparing cell-position projections onto 360 equally spaced directions. The confinement measure, 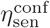, was the minimum number of sensitive cells lying at or beyond the outermost resistant cell across these directions, normalized by the initial total cell number. The escape measure, 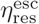, was the number of unique resistant cells extending beyond the sensitive population in at least one direction, normalized by the same initial total. TTE was the first observation with 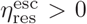. Treatment failure was defined as the first observation at which the resistant population reached 50% of the initial total population. The tracking procedure, the complete mathematical definitions of the confinement and escape measures as well as the associated inclusion criteria are given in Supplementary Methods Section 2. Local total, sensitive and resistant cell-density fields were reconstructed from cell coordinates using an Epanechnikov kernel estimator with a radius of 120 *μ*m. Resistant growth was then related to local total density between consecutive observations. This analysis tests whether the local crowding coordinate associated with the spatial model also organizes the observed resistant-growth dynamics. Details of point filtering, grid resolution, kernel normalization, interval selection and binning are given in Supplementary Methods Section 3.

### Spatial particle-based model

To provide a mechanistic representation of the experimental system, we employed a particle-based model based on dissipative particle dynamics, building on previously established tissue-mechanics frameworks [23, 26, 27]. Individual cells were represented by pairs of mechanically interacting particles and underwent growth, division and stochastic death, thereby capturing the spatial structure of the experimental colonies.

Cell expansion was modulated by local mechanical constraints. Together with a size-dependent division probability, this resulted in crowding-mediated proliferation rate.

Therapy was modeled using first-order phenomenological equations describing the time-dependent environmental concentration, including rapid uptake during treatment and slower decay in its absence (Supplementary Methods Section 4.1). Cellular drug concentrations relax towards the environmental concentration on a characteristic response timescale, introducing a delay between treatment and cellular response. Therapy-sensitive cells undergo stochastic apoptosis with a death rate proportional to their intracellular drug concentration and cell size, whereas resistant cells are unaffected. The model therefore captures the key mechanisms investigated experimentally: spatial competition, crowding-mediated growth suppression, resistance and delayed treatment response.

Simulations were initialized from experimentally measured colony configurations and parameterized using experimentally observed growth and treatment-response dynamics. Growth-related parameters were first constrained using untreated colonies, after which treatment-response parameters were inferred from continuously treated and adaptive-therapy experiments. Parameter estimation was performed by comparing simulated and experimental colony growth, cell numbers and spatial density distributions. A single parameter set was used across all treatment protocols without protocol-specific tuning.

Additional details of the model formulation and parameter-inference procedure are provided in Supplementary Methods Section 4. The complete simulation parameter set is listed in Supplementary Table S1.

### Effective model and parameter calibration

To obtain a computationally efficient representation of pre-escape dynamics, we used a two-population generalized Lotka–Volterra model for normalized sensitive (*S*) and resistant (*R*) cell numbers:

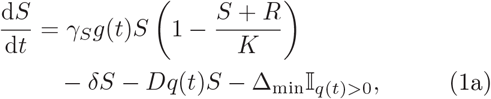

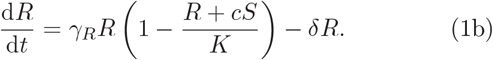

Here, *γ*_*S*_ and *γ*_*R*_ are the population growth rates, *K* is the carrying capacity, *δ* is the baseline death rate, *D* and Δ_min_ describe treatment-associated sensitive-cell killing, and *c* quantifies the competitive effect of sensitive cells on resistant cells. The functions *g*(*t*) and *q*(*t*) account for delayed suppression of sensitive-cell growth and induction of treatment-associated death after treatment initiation and withdrawal. Resistant cells were not directly affected by treatment. Numerical implementation and parameter values are described in Supplementary Methods Section 5.1.

Parameters were fitted to 32 experimental trajectories, comprising eight replicates each from no-treatment, single-pulse, CT and AT100 conditions. Cell numbers were normalized to the combined sensitive and resistant count at the first observation. The first up to nine available observations per trajectory were retained, yielding 266 observation times, each contributing sensitive and resistant counts. Each simulation was initialized using the observed sensitive and resistant fractions of the corresponding trajectory.

Treatment was fixed to off and on for no-treatment and CT trajectories, respectively, and was applied only during the first interval for pulse-treated trajectories. For AT100, the action sequence was reconstructed from the observed experimental burden, with treatment applied when the observed normalized burden was greater than or equal to its initial value. AT100 was therefore included in calibration.

For trajectory *e* with *T*_*e*_ observations, the calibration loss was the mean, across the *E* = 32 trajectories, of the sum of the sensitive- and resistant-population mean abso-lute errors,

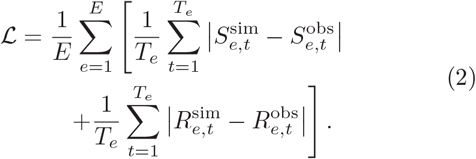

Uncertainty was assessed by resampling complete trajectories with replacement within each treatment condition. We generated 20 stratified bootstrap datasets, each containing eight trajectories per condition, and refitted the model from the frozen reference parameter vector. All 20 terminal parameter vectors were retained and used to train one RL agent each; optimizer outcomes and random seeds are reported in Supplementary Methods Section 5.1.

### Reinforcement-learning environment and training

Treatment policies were trained using Proximal Policy Optimization implemented in Stable-Baselines3 [45]. At each decision point, the agent selected treatment off or on for the following interval according to the experimental 2–2– 3-day schedule.

The policy input comprised the four most recent pairs of normalized total burden N = S +R and treatment state u (0=OFF or 1=ON):

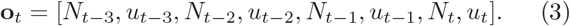

The agent had no access to clonal composition, spatial information, local density, calendar time or model parameters. In the reconstructed training pipeline, episodes began at *S*_0_ = 0.9995 and *R*_0_ = 0.0005 and ended when *N*≥3*N*_0_ or after 60 decisions, corresponding to 140 days.

Policies comprised separate actor and critic networks with two hidden layers of 64 units each. Training proceeded in two stages. During pretraining, a reward of +1 was assigned when treatment was withheld and N < 1.5N0; all other transitions received zero reward. During fine-tuning, each transition before termination or truncation received +1, thereby optimizing episode duration. The requested training lengths were 100,000 and 160,000 environment steps, respectively. Checkpoints were selected using the running mean reward of recent training episodes, not an independent validation environment.

The preserved Proximal Policy Optimization checkpoint used to generate the in vitro ATagent schedule was retained as the experimental reference policy. Separately, the 20 bootstrap-derived parameterizations were each used to train one Proximal Policy Optimization agent in a standardized retrospective workflow; their deterministic decisions are summarized in Fig. 4d. Implementation hyperparameters and the provenance of the preserved experimental checkpoint are reported in Supplementary Methods Section 5.2.

### Policy-behaviour analysis

To characterize the behaviour of the trained agents, we directly evaluated their deterministic treatment decisions across a predefined synthetic observation space matching the four-frame input used during training.

Synthetic burden histories were constructed from a baseline burden *b* by adding a single peak of magnitude Δ*p* at position *d ∈ {* 0, 1, 2, 3 *}*, ordered from oldest to most recent observation. All 16 possible four-action treatment histories were evaluated. Baseline burdens and peak magnitudes ranged from 0 to 5 in increments of 0.5. Only burden histories satisfying *b* + Δ*p≤* 3 were retained, ensuring that all inputs remained within the trained observation range without clipping.

For visualization, histories were summarized by the growth score

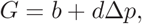

which assigns greater weight to peaks occurring in more recent observations. Growth scores were assigned to 14 equal-width bins spanning 0–9. Within each agent, growth-score bin and exact treatment history, the treatment-on fraction was calculated as the fraction of deterministic actions equal to treatment on. Figure 4d shows the mean and sample standard deviation across the 20 bootstrap-trained agents for the five histories [0, 0, 0, 0], [0, 0, 0, 1], [0, 0, 1, 1], [0, 1, 1, 1], and [1, 1, 1, 1]. The analysis therefore maps how deterministic treatment decisions depend jointly on burden dynamics and recent treatment history. Exact observation counts and policy exclusions are reported in Supplementary Methods Section 5.3.

### Statistical analysis

Independent experimental units were individual colonies and independent computational units were simulation runs. Time-to-event outcomes were analysed using Kaplan–Meier estimators, with trajectories without an observed event right-censored at their last valid observation. Groups were compared using two-sided log-rank tests; for the all-pairwise survival comparisons in Fig. 4h and i, p values were Bonferroni corrected. Effect sizes are reported as median differences with 95% percentile-bootstrap confidence intervals where applicable. For analyses of repeated observations, resampling was performed at the colony level, retaining all available time points from each sampled colony; repeated observations were otherwise used for descriptive trajectory summaries and were not treated as independent biological replicates. Spatial density curves are descriptive pooled-pixel summaries; biological replication is represented by colonies or simulation runs. Exact sample sizes, event counts, statistical tests and definitions of error bars are reported in the relevant figure legends. Replicate-level exclusions and analysis cutoffs are documented in Supplementary Methods Section 6 and Supplementary Table S3.

## Data availability

Raw imaging datasets are too large for public deposition but are available from the corresponding author upon reasonable request.

## Code availability

Custom code used for the simulations is available at https://github.com/KayserLab/Eiche_Fischer_2026.git. The repository contains all scripts necessary to reproduce the simulation results and figures presented in this study.

## Acknowledgements

The authors thank N. Appold, I. Schneider, L. von Selzam and A. Palm for valuable discussions. We acknowledge the contribution of L. Nuić in the code development. We are extremely grateful for the support of J. Guck and his entire division.

## Author Declaration

ME, EF, and JK conceived of the overall study design for experiments. MM and ASS conceived of and designed the DPD simulations. ME constructed the cell lines. EF developed the experimental assay and the same-day analysis and decision pipeline. MM developed the digital twin platform. ME and EF conducted experiments and image analysis. MM performed and analyzed DPD simulations. SA, TC, ME, and JK built the abstraction and reinforcement learning pipeline. MM and AS performed agent behavior analysis. ME, EF, and MM jointly analyzed all data. JK and ASS provided supervision and oversaw the execution of the project. ME, EF, MM, ASS, and JK wrote the manuscript.

