## Supplementary Material for "Crowding governs resistance confinement and guides the improvement of adaptive therapy in a 2D melanoma model system"

### Supplementary Information

**Supplementary Figures** **8**

**Supplementary Tables** **16**

### Supplementary Methods

#### 1 Image segmentation and saturation correction

Raw CZI images were processed independently for the sensitive and resistant fluorescence channels. Cell masks were generated using the custom Cellpose v3.0.11 models `a4_cyto2_PhD_1000_new` for the resistant channel and `a2_cyto3_Epi_1000_PhD` for the sensitive channel [33]. The frozen model files and their SHA-256 checksums are provided in `analysis/cellpose/models/` of the manuscript repository.

The resistant-channel model was trained using 39 maximum-intensity-projection or cropped fluorescence image–mask pairs from G418 b2b4 and a2a4 experiments acquired between November 2023 and April 2025; 38 pairs retain Cellpose manual-edit records. The sensitive-channel model was trained using 19 epifluorescence maximum-intensity-projection or cropped image–mask pairs from G418 a2a4 and b2b4 experiments acquired between February 2024 and March 2025; all 19 pairs retain manual-edit records.

Segmentation used two-dimensional `CellposeModel.eval` with `diameter=None`. Consequently, the stored model-specific mean training-mask diameters of 31.24 and 23.87 pixels were used for resistant and sensitive cells, respectively, rather than per-image automatic diameter estimation. A minimum object size of 250 pixels, a flow threshold of 0, a cell-probability threshold of $-1$, and a stitch threshold of 0 were used. The centroid of each accepted mask was used as the corresponding cell coordinate. Images were processed without background subtraction or interchannel bleed-through correction.

To identify regions in which cell numbers were underestimated because of detector saturation, pixels with intensities above 12,000 that were not assigned to accepted cell masks were identified. The number of missing cells per image channel was estimated by multiplying their total area by a calibration factor of 0.00286 cells per pixel. For spatial analyses, reconstructed coordinates were generated by Poisson-disc sampling globally over the cleaned saturation mask. The initial sampling separation was 15 pixels and was relaxed to a minimum of 4 pixels when necessary; the sampling seed was fixed for reproducibility. Original and reconstructed coordinate sets were centre-padded to a common image canvas before mask-based analyses.

#### 2 Resistant-clone tracking and confinement and escape calculation

In experiments, resistant masks were morphologically closed using a disk with a radius of three pixels, after which holes were filled. Initial resistant components were grouped by their centroids using DBSCAN with a 200-pixel neighbourhood and a minimum of one component per cluster. Components were linked between consecutive observations by direct overlap or, when necessary, by overlap after label expansion over a distance of 150 pixels multiplied by the number of elapsed days. When tracked components merged, the identity associated with the largest preceding tracked-mask area was retained. Sensitive-cell coordinates were rasterized into occupancy masks, morphologically closed using the same three-pixel disk, and subjected to hole filling. Sensitive components were retained if they overlapped the preceding retained sensitive mask, directly or after expansion using the same daily linking distance. When the preceding retained sensitive mask was absent or empty, all occupied components were retained.

A resistant clone was included in the escape analysis only if at least ten resistant-cell coordinates were assigned to it at its last observation containing one or more mapped resistant-cell coordinates. Resistant coordinates were retained only within eligible tracked-clone masks, and sensitive coordinates were retained only within the cleaned sensitive masks. All eligible resistant clones contributed jointly to the geometric calculation.

Confinement and escape were evaluated using cell-position projections onto 360 equally spaced directions. For $\theta_k = 2\pi k/360$, where $k \in \{0, \ldots, 359\}$, let $\mathbf{u}_{\theta_k} = (\cos\theta_k, \sin\theta_k)$ denote the corresponding unit vector and let

$$p_i(\theta_k) = \mathbf{x}_i \cdot \mathbf{u}_{\theta_k} \tag{4}$$

denote the projection of the position $\mathbf{x}_i$ of cell $i$ onto this direction. For observations containing both retained sensitive and resistant cells, the directional confinement count was defined as

$$c_{\theta_k} = \#\left\{s : \; p_s(\theta_k) \geq \max_r p_r(\theta_k) - \varepsilon\right\}, \tag{5}$$

where $s$ and $r$ denote retained sensitive and resistant cells, respectively, and $\varepsilon = 10^{-9}$ coordinate units is the numerical projection tolerance. The set of resistant cells extending beyond the sensitive population in at least one direction was

$$E = \bigcup_{k=0}^{359} \left\{r : \; p_r(\theta_k) > \max_s p_s(\theta_k) + \varepsilon\right\}. \tag{6}$$

Each resistant cell was counted only once, irrespective of the number of directions in which it extended beyond the sensitive population. The normalized confinement and escape measures were defined as

$$\eta_{\text{sen}}^{\text{conf}} = \frac{\min_{k} c_{\theta_k}}{N_0}, \tag{7}$$

and

$$\eta_{\text{res}}^{\text{esc}} = \frac{|E|}{N_0}, \tag{8}$$

respectively. Thus, the confinement measure quantifies the smallest sensitive-cell count lying at or beyond the resistant front across the sampled directions, whereas the escape measure quantifies resistant cells extending beyond the sensitive population. Both measures were non-negative and were recalculated independently at each observation.

Here, $N_0$ denotes the initial total cell number used to normalize the corresponding plot. Experimental escape trajectories displayed alongside whole-colony population burdens used the initial total from the population-count input. Supplementary escape time courses used the initial retained sensitive- and resistant-cell total; initial-confinement histograms used this total at the first valid observation. Simulated escape trajectories were normalized using the initial retained total cell count.

For observations with resistant cells but no sensitive cells, confinement was set to zero and all retained resistant cells were counted as escaped. Observations without retained resistant cells had an undefined confinement count and an escaped-resistant count of zero. Such observations were omitted from TTE determination; the undefined confinement count was displayed as zero in the supplementary escape time courses. Replicates without any valid resistant-cell observations did not contribute a TTE.

Time to escape (TTE) was defined as the first sampled acquisition day satisfying $\eta_{\text{res}}^{\text{esc}} > 0$, with the first observation designated day 1. Experimental replicate exclusions and follow-up cutoffs were applied according to the predefined quality-control rules. Simulation TTE analyses included only runs with numerical identifiers greater than 9.

#### 3 Local-density estimation and master-curve construction

Cell densities were evaluated on a regular grid with a spacing of approximately 26.1 $\mu$m. Local density fields were obtained using a normalized circular Epanechnikov kernel of radius $R = 120$ $\mu$m,

$$K_R(r) = \frac{2}{\pi R^2}\left(1 - \frac{r^2}{R^2}\right)\mathbb{I}(r \le R), \tag{9}$$

where $\mathbb{I}$ denotes the indicator function.

Before density estimation, experimental wells were excluded or truncated according to the predefined experimental quality-control rules. No additional spatial outlier filter or recentering was applied to experimental coordinates. Simulated coordinates were filtered using radial and local-neighbour criteria. Cells located more than 504.8 simulation units from the origin were discarded, and retained cells were required to have at least five cells, including the focal cell, within 30.3 simulation units. Under the simulation-to-experiment calibration, these thresholds correspond to 6,000 and 360 experimental pixels, respectively.

For each retained interval from $t_0$ to $t_1$, the density coordinate was defined as the mean total density at the two interval endpoints,

$$\rho_{\text{tot}} = \frac{\rho_{\text{tot}}(t_0) + \rho_{\text{tot}}(t_1)}{2}. \tag{10}$$

The corresponding relative change in resistant-cell density per unit time was

$$\frac{\Delta\tilde{\rho}_{\text{res}}}{\Delta t} = \frac{\rho_{\text{res}}(t_1) - \rho_{\text{res}}(t_0)}{\rho_{\text{res}}(t_0)(t_1 - t_0)}. \tag{11}$$

For the all-timepoints curves, all consecutive intervals were retained after any well-specific truncation. Simulation intervals were 24 h long, whereas experimental changes were divided by the actual elapsed time between acquisitions, which was typically two or three days. Grid-point–interval observations were retained only where at least 14 cells were located within the kernel radius at both endpoints. They were additionally required to satisfy

$$\rho_{\text{res}}(t_0) \ge \frac{2.25}{\pi R^2} \tag{12}$$

at the initial endpoint. Simulation master curves included only runs with numerical identifiers greater than 9.

Within each run or well, the retained grid-point–interval observations were pooled and grouped into total-density bins with a width of approximately 0.063 cells/1,000 $\mu m^2$. Bins containing fewer than 20 observations were omitted, and the median value was calculated for each remaining bin. Each strategy curve shows the median across the resulting run- or well-level curves, with the interquartile range shown as shading. At least three valid run or well curves were required in each bin. The master curve is the mean of the strategy curves, and its shaded region spans their minimum to maximum values. At least two valid strategies were required in each master-curve bin.

### 4 Spatial particle-based model

#### 4.1 Model description

To obtain a mechanistically explicit description of tissue dynamics, we employ simulations based on dissipative particle dynamics (DPD, Fig. 1a) [23]. Our implementation builds on the framework of [26, 27], in which each cell is modeled as a pair of particles with a well-defined cell size. Cells interact via short-range conservative, dissipative, and stochastic forces, including intracellular growth forces driving particle separation, intercellular repulsion and adhesion, and pairwise dissipative interactions, with functional forms following [27]. Cell shapes and areas are approximated using a center-of-mass Voronoi tessellation constructed from nuclei positions (Fig. 1b).

To capture the mechanical influence of the environment on the cell life cycle, population dynamics are governed by cell-size-regulated stochastic rules for division and apoptosis, described in [27].

The effect of therapy is incorporated through a coarse-grained description of drug concentration dynamics (Fig. 1c). We introduce a homogeneous environmental therapy concentration $C(t)$ and a cell-specific internal concentration $c_i(t)$. Both quantities represent dimensionless exposure levels normalized to the interval $[0, 1]$, where 0 corresponds to the absence of therapy and 1 to full saturation.

The temporal evolution of $C(t)$ is governed by a binary switch function $\sigma(t) \in \{0, 1\}$ representing intermittent drug administration matched to the experimental treatment schedule. The dynamics of $C(t)$ are given by a delayed-response equation

$$\frac{dC(t)}{dt} = \begin{cases} \dfrac{1 - C(t)}{\tau_{\mathrm{abs}}}, & \sigma(t) = 1 \\ -\dfrac{C(t)}{\tau_{\mathrm{red}}}, & \sigma(t) = 0 \end{cases} \tag{13}$$

where $\tau_{\mathrm{abs}}$ and $\tau_{\mathrm{red}}$ denote the characteristic absorption and decay timescales, respectively. We consider the regime $\tau_{\mathrm{abs}} \ll \tau_{\mathrm{red}}$, such that the environmental concentration rapidly approaches saturation during treatment periods and slowly relaxes in the absence of treatment due to cellular uptake and degradation.

At the single-cell level, the internal concentration $c_i(t)$ relaxes toward the environmental concentration

$$\frac{dc_i(t)}{dt} = \frac{C(t) - c_i(t)}{\tau_{\mathrm{resp}}}, \tag{14}$$

where $\tau_{\mathrm{resp}}$ is the characteristic cellular response timescale. Upon cell division, daughter cells inherit the concentration of the parent cell.

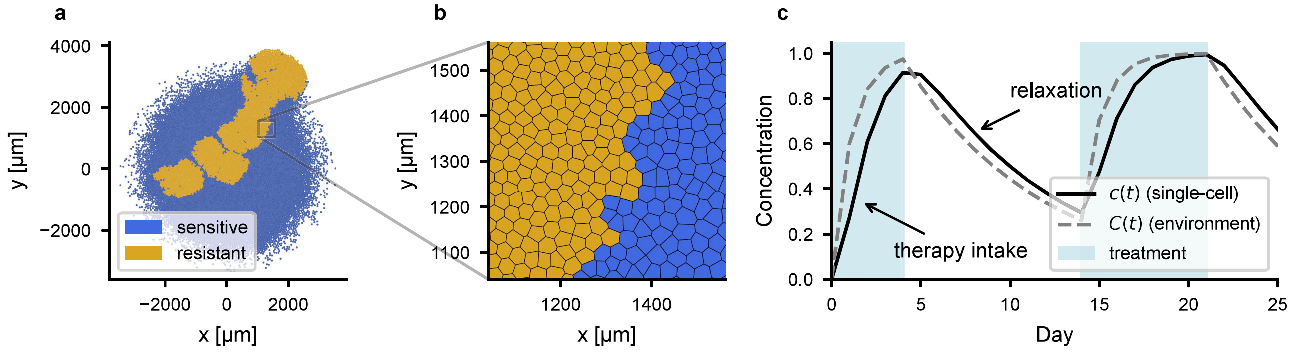

Supplementary Figure 1: ***In silico* model of melanoma colonies.** **a** A snapshot of a simulated tissue (AT100, day 44) showing sensitive and resistant cells. **b** Zoomed in view of panel **a**. Cell membranes are calculated as the Voronoi tesselation of cell centers. **b** Time evolution of environment and single-cell therapy concentration levels described by equations (13) and (14). Shaded regions mark the treatment schedule defined by the protocol.

To model the effect of therapy on cell viability, we introduce a concentration-dependent apoptosis rate with a weak cell-size dependence (motivated by experimental observations).

Specifically, the therapy-induced death rate of cell $i$ is defined as

$$k^i_{\text{death}} = \Phi\sqrt{\frac{A_i}{A_0}}\, c_i(t), \tag{15}$$

where $A_i$ is the instantaneous cell area, $A_0$ is a reference area corresponding to the maximal cell size, and $\Phi$ sets the overall strength of the therapy effect.

The survival probability over a timestep $\Delta t$ is therefore modeled as a Poisson process ($P^i_{\text{survive}} = \exp\left(-k^i_{\text{death}}\Delta t\right)$.

We distinguish between therapy-sensitive and therapy-resistant cell populations through their response to this process. Both populations accumulate the internal concentration $c_i(t)$, but only sensitive cells undergo therapy-induced apoptosis, while resistant cells are unaffected.

##### 4.2 Initialization and parameter inference

Simulations are initialized using experimentally measured nuclei positions of sensitive and resistant cells at day 1 for each protocol (Fig. 1h,j,l).

We adopt a baseline set of mechanical parameters from [27], previously calibrated for epithelial tissues. The mapping between simulation and experimental units is established by matching the characteristic cell size and the doubling time measured in experiments. All simulation outputs are rescaled accordingly before comparison to data. In these units, one simulation timestep corresponds to $10^{-5}$ days. To scale the simulation data to the experiments, we take $S = \sqrt{A_{\text{exp}}/A_{\text{sim}}}$, where $A_{\text{exp}}$ ($A_{\text{sim}}$) is the cell area of the leading-edge susceptible cells observed in experiment (simulation).

To parametrize the system, we started from a previously calibrated epithelial tissue model [27], under the assumption that melanoma cells exhibit comparable mechanical behavior and that macroscopic dynamics are insensitive to microscopic force details. Further parameter tuning was performed in a hierarchical manner (first for the untreated colonies, then for continuous therapy, and finally for AT) through a multiparameter fit using a Globalized Nelder–Mead algorithm [46] and manual fitting. Simulations are compared to experiments via the temporal evolution of type-dependent normalized cell numbers, and the resulting fits are shown in Fig. 1i,k,m.

A single consistent parameter set is used across all treatment protocols, enabling direct comparison between simulations and experiments without protocol-specific tuning. The full set of fitted parameters is summarized in Table S1.

##### 4.3 Simulation database and protocol validation

To enable systematic comparison across treatment protocols, we constructed a database of initial conditions derived from both experimental data and synthetic configurations. The dataset comprises 40 experimentally obtained initial conditions, corresponding to approximately 10 samples per protocol (CT, AT50, AT100, and AT$^+$), and 10 additional synthetic initial conditions generated from a single experiment. The synthetic configurations were created by selecting a region of sensitive cells and introducing a localized cluster of mutant cells at varying distances from the colony center.

Each of the 50 initial conditions was then simulated under all five treatment protocols (NT, CT, AT50, AT100, and AT$^+$), as well as under policies generated by the reinforcement learning (RL) agent. Simulations were run until either tumor progression was reached or numerical instabilities caused termination. Simulations that did not reach progression were retained as right-censored observations at their final available output time. Figure 4 comparisons excluded synthetic initial conditions and used available experiment-derived runs: CT 36, AT50 35, AT100 36, ATagent 36, and AT$^+$ 24.

##### 4.4 Mechanoregulation via local density

Growth requires a cell to increase its size against forces exerted by its neighbours (Fig. 3a), thus setting the pace of single-cell expansion. We test this directly with individual-cell data from the simulation: for every cell that survives a $\Delta t = 12\,\text{h}$ window without dividing or dying, we compute the expansion rate $\lambda = (d(t+\Delta t) - d(t))/\Delta t$, where $d$ is the distance between the two particles composing the cell. Resolving $\lambda$ against the local density $\rho_{\text{tot}}$ around each cell (Fig. 3c) shows that expansion is fastest in sparse regions and is progressively suppressed as $\rho_{\text{tot}}$ increases, for both susceptible and resistant cells.

#### 5 Effective-model implementation and reinforcement learning

##### 5.1 Numerical implementation and fitting details

The effective model was integrated using an explicit Euler-type update with $\mathrm{d}t = 0.0025$ in internal model-time units. Two-day decision intervals comprised 1,600 integration steps, corresponding to four internal time units, whereas three-day intervals comprised 2,400 steps, corresponding to six internal time units.

| Parameter | Value | Description |
|---|---|---|
| $\mu_{\text{bg}}$ | 0.154 | Substrate dissipation |
| | | *Therapy* |
| $\tau_{\text{abs}}$ | 1.086 | Therapy absorption timescale |
| $\tau_{\text{red}}$ | 7.6 | Therapy reduction timescale |
| $\tau_{\text{resp}}$ | 0.86 | Cellular response timescale |
| $\Phi$ | 1.17 | Therapy efficacy coefficient |
| | | *Growth* |
| $B, B^{\text{mut}}$ | 1.9, 2.4 | Growth force amplitude (wild-type, mutant) |
| $r_0$ | 1.12 | Regularization length preventing divergence of growth force |
| $A_0$ | 18.34 | Bounding cell area |
| | | *Dissipation and interactions* |
| $\gamma_c$ | 1.1 | Intracellular dissipative coefficient |
| $R_t$ | 2.5 | Interaction range of dissipative forces |
| $\mu_{\parallel}$ | 0.2 | Intercellular dissipation along connection-line |
| $\mu_{\perp}$ | 0.15 | Intercellular dissipation perpendicular to connection-line |
| $f_0$ | 150.0 | Strength of short-range repulsion |
| $f_1, f_1^{\text{mut}}$ | 3.64, 2.32 | Adhesion strength (wild-type, mutant) |
| $R_{pp}$ | 1.83 | Range of pairwise interaction potential |
| $\{$ | 1.4 | Scaling factor for interaction cutoff and neighbor search |
| | | *Division and death* |
| $k_{\text{apoptosis}}$ | 0.001 | Regular apoptosis rate |
| $R_{c1}, R_{c1}^{\text{mut}}$ | 1.5, 2.24 | Lower size threshold for division (wild-type, mutant) |
| $R_{c2}, R_{c2}^{\text{mut}}$ | $(1.1, 2.6) \times 10^4$ | Upper size threshold for division (wild-type, mutant) |
| $r_c$ | 0.01 | Initial separation of daughter particles after division |
| | | *Noise and additional terms* |
| $k_B T$ | 0.424 | Effective noise strength |
| $m$ | $5.76 \times 10^{-4}$ | Effective particle mass |
| $f_3$ | 1.0 | Density-gradient force coefficient |
| $\tau_{\text{div}}$ | 0.25 | Duration of division phase |
| $K_s$ | 1000.0 | Spring constant during division |
| $\lambda_{\text{Lloyd}}$ | $5 \times 10^{-6}$ | Strength of Lloyd relaxation |

Table S1: Model parameters used in the simulations. The timestep is $10^{-5}$ days.

The treatment response of sensitive cells was represented by separate internal growth- and death-response fractions. Following treatment initiation, growth suppression and treatment-associated death commenced after an onset delay and changed gradually. Following treatment withdrawal, the treatment state persisted for an offset delay before recovery. The minimum killing term was applied only while the treatment-death fraction was positive. Treatment-induced death of resistant cells was set to zero.

The fixed reference values retained with the reconstructed RL workflow were $\gamma_S = 0.081$, $\gamma_R = 0.223$, $c = 5.21$, $K = 7.87$, $\delta = 2 \times 10^{-5}$, $D = 0.0975$, and $\Delta_{\text{min}} = 0.011$. The treatment-response ramp-up and ramp-down parameters were 13.8 and 0.1, respectively, and the treatment-on and treatment-off delays were 8.7 and 10.6. Values are reported in the internal normalized units of the implementation because a physical-unit mapping for all historical time constants was not retained.

The calibration dataset contained eight trajectories per treatment condition. Twenty-four trajectories contributed nine observations, two contributed seven observations, and six contributed six observations; all shortened trajectories belonged to the no-treatment condition. The first recorded measurement, corresponding to day 1 in the archived data, defined the trajectory-specific normalization and initial condition.

Eleven optimizer coordinates were varied, but ($\Delta_{\text{min}}$) remained fixed at 0.011 during fitting. Its returned coordinate, which did not affect fitting loss, was subsequently applied during bootstrap-agent training. Parameter initialization and bounds are provided with the executable fitting workflow. The fixed parameter vector preserved in the RL directory was used as the common initialization for the retrospective bootstrap fits. Although the current workflow reproducibly refits the model from the experimental trajectories, the exact historical fitting run that generated this archived parameter vector could not be reconstructed.

Of the 20 bootstrap fits, 14 terminated with optimizer success and six reached the iteration limit. All 20 terminal parameter vectors were retained. One PPO agent was trained per parameter vector using seed $20260600 + i$, where $i = 0, \dots, 19$. Five fixed-parameter repeated-seed agents and one control agent were generated for diagnostic purposes but excluded from Fig. 4f.

##### 5.2 Reinforcement-learning implementation and checkpoint provenance

The bootstrap agents were trained using Stable-Baselines3 version 2.2.1 with `MlpPolicy`, separate [64, 64] actor and critic networks, a rollout length of 64, a batch size of 32, four optimization epochs per update, a learning rate of $10^{-4}$, an entropy coefficient of $10^{-5}$, a clip range of 0.005, a discount factor of 0.99, and a generalized-advantage-estimation parameter of 0.95. The checkpoint callback retained models according to the running mean reward over the eight most recently completed training episodes.

The historical checkpoint used to generate the *in vitro* ATagent schedules is preserved in the manuscript repository. Its stored final-stage metadata specify [64, 64] actor and critic networks, a rollout length of 128, a batch size of 64, eight optimization epochs per update, a learning rate of $10^{-4}$, an entropy coefficient of $10^{-5}$, and a clip range of 0.005. The checkpoint was saved after 119,936 fine-tuning timesteps; no explicit PPO seed was stored.

The original effective-model environment and complete stage-one training configuration used to generate the historical checkpoint are unavailable. Accordingly, the preserved checkpoint itself, rather than a reconstructed approximation, was used for the biological experiments. The 20 bootstrap-trained policies were generated separately in the standardized retrospective workflow described above.

##### 5.3 Synthetic policy probing

To characterize the treatment policies learned by the trained agents, we evaluated them on a systematically constructed synthetic observation space, preserving the four-observation structure used during training. Agents use normalized total tumor burden and therapy decisions on observation days 0–3 while allowing the policy response to be examined under controlled conditions. This construction (Methods) generated 7,744 candidate observations per agent. Restricting to burden histories satisfying $b + \Delta \leq 3$ retained 1,792 in-domain observations per agent, none of which required clipping.

We considered all 16 possible treatment histories, binary sequences from 0000 to 1111 indicating whether treatment was on or off at each observation day. For the burden observations we did a linear-response analysis. One day was chosen to be the peak day and assigned the highest normalized burden, while the remaining three days shared a common base value. Burden trajectories were constructed by varying this base value and the additional burden at the peak day.

Each synthetic sequence of four burdens and four treatment decisions was passed through the trained agents. We summarized the burden trajectory using a history growth score, defined in the main text, and computed the treatment-on fraction as the fraction of decisions that initiated or continued treatment.

Agents were more likely to treat when the growth score was high, particularly when the burden increase occurred at a recent observation and treatment had not been given in the preceding days, consistent with short, intermittent treatment episodes rather than sustained treatment.

To assess robustness, we repeated this analysis across the ensemble of bootstrap-trained agents and across repeated-seed training runs. For each agent, deterministic actions were averaged within each exact treatment history and populated growth-score bin to obtain a treatment-on fraction. Figure 4f reports bootstrap-agent mean and sample standard deviation for histories pooled by latest action; dashed curves show the historical experimental agent. Repeated-seed and control agents are excluded from the bootstrap summary.

#### 6 Replicate exclusions and analysis cutoffs

Supplementary Table S3 lists all replicate-level complete exclusions and endpoint cutoffs. Complete exclusions were applied when resistant cells were absent or when a replicate could not contribute interpretable data. For otherwise usable replicates, an endpoint cutoff was applied when loss of resistant cells from the imaged colony rendered subsequent population counts non-representative. This cutoff typically corresponded to the loss of approximately 20% of the resistant-cell population. Only observations up to and including the listed final retained day were included in subsequent analyses.

#### Supplementary Figures

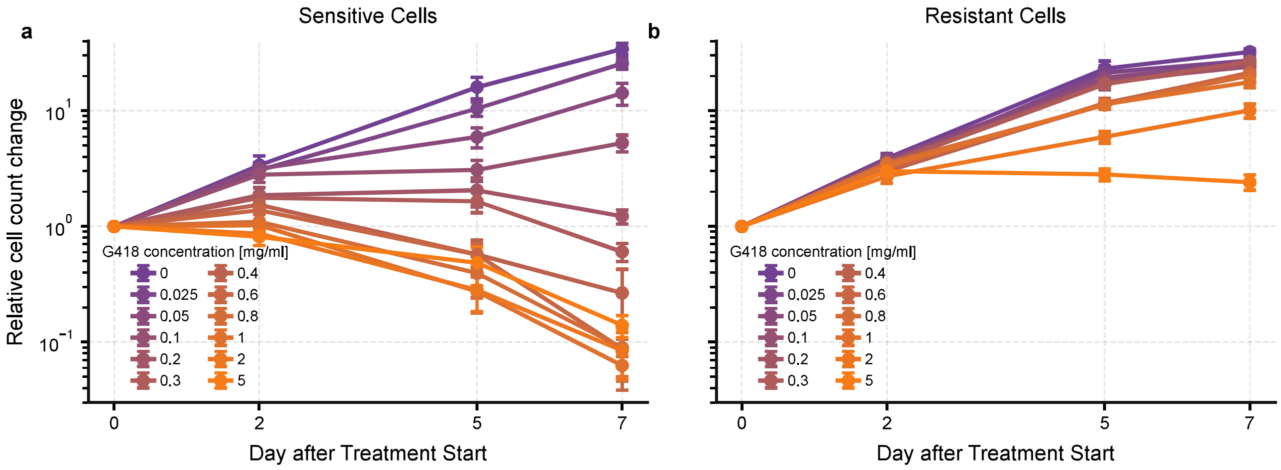

Supplementary Figure 2: **Differential growth and drug-response dynamics of $\alpha 2$ and $\alpha 4$ melanoma cell lines under G418 treatment.** Panels **a** and **b** show the growth responses of $\alpha 2$ and $\alpha 4$ cells, respectively, across increasing G418 concentrations over seven days. Metabolic activity was quantified using the AlamarBlue assay described in the Methods and normalized to the pretreatment measurement. Data points show the mean relative change from pretreatment, and error bars indicate the SD across eight technical replicates per condition.

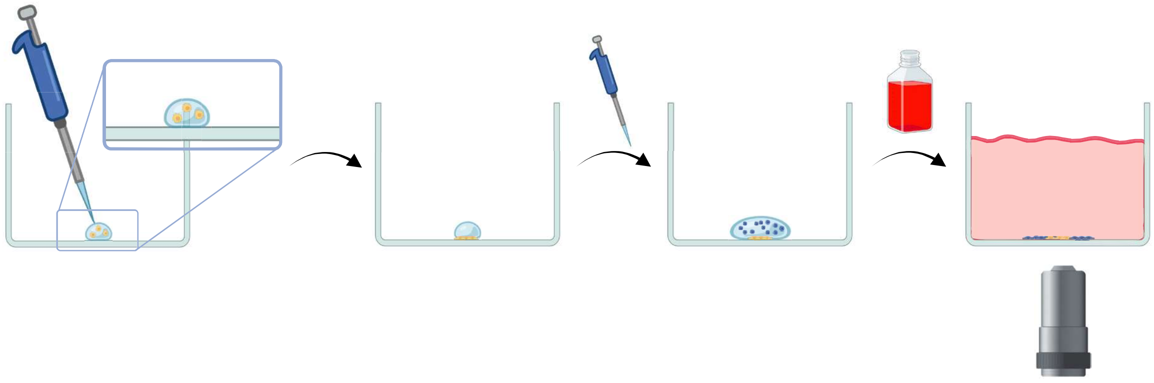

Supplementary Figure 3: **Detailed sequential seeding workflow.** Resistant cells (yellow) were first seeded centrally as a 3 $\mu$L droplet at 1,333 cells/ml and allowed to adhere for 4–5 h. A 20 $\mu$L droplet of sensitive cells (blue; $1 \times 10^6$ cells/ml) was subsequently added at the same position and allowed to adhere for a further 18 h. RPMI medium supplemented with 10% FBS was then carefully added to cover the well, yielding radially confined colonies with centrally positioned resistant cells surrounded by sensitive cells. The inset illustrates the initial placement of resistant cells within the seeding droplet. Created with BioRender.com.

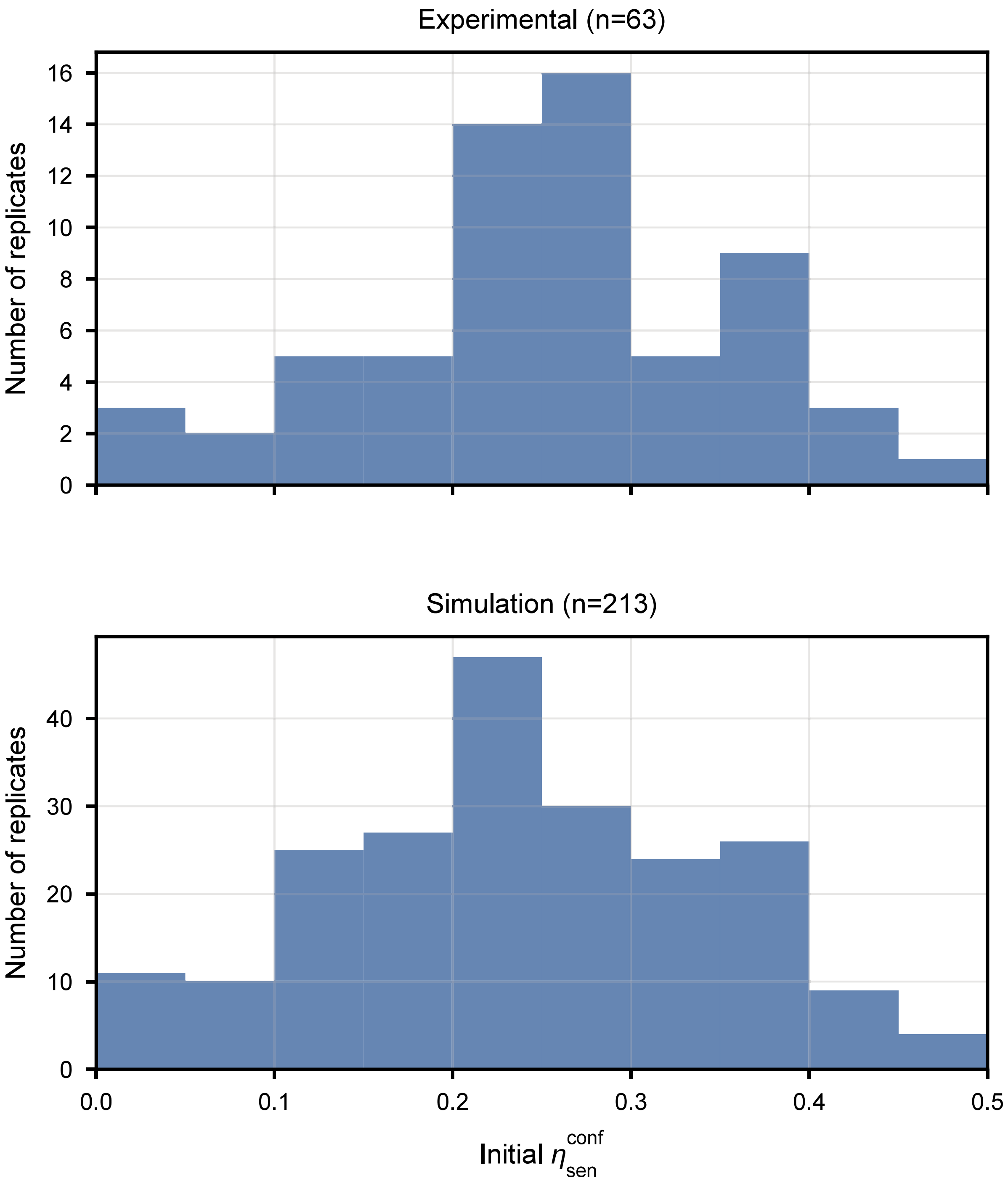

Supplementary Figure 4: **Distribution of the initial escape margin across experimental and simulated replicates.** Histograms show the initial normalized confinement margin, $\eta_{sen}^{conf}$, for experimental replicates (top; $n = 63$) and simulated replicates (bottom; $n = 213$).

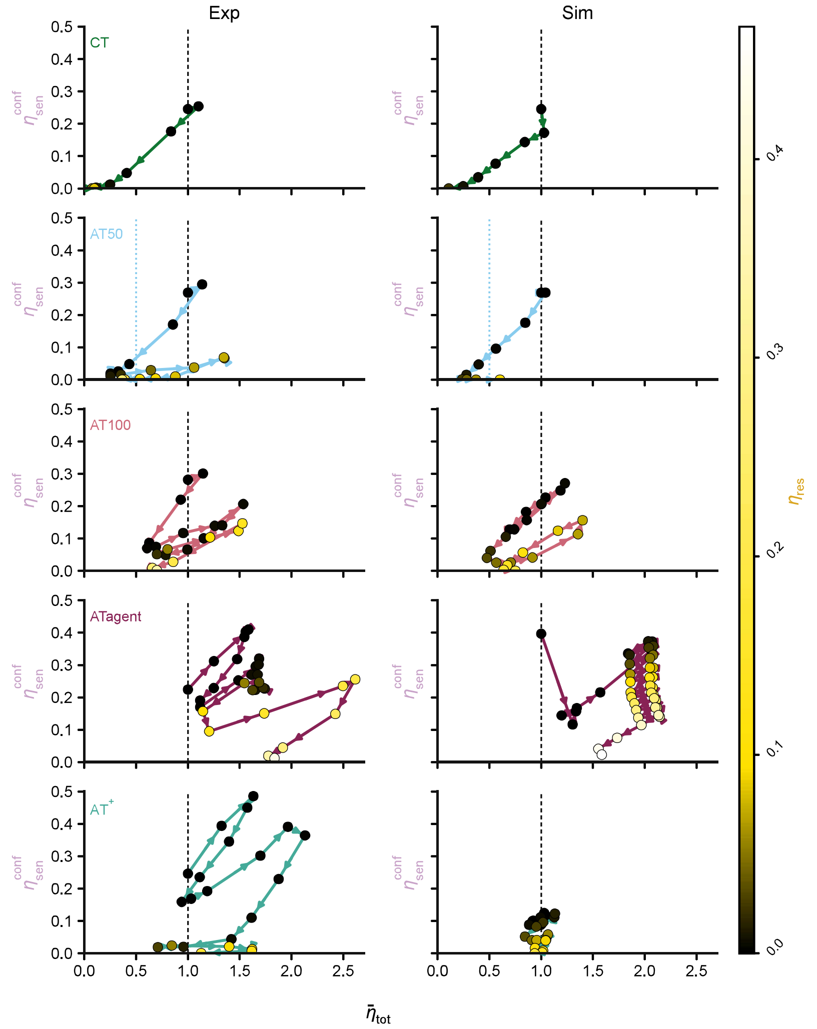

Supplementary Figure 5: **Confinement margin as a function of total cell burden across treatment strategies.** Confinement margin $\eta_{\mathrm{sen}}^{\mathrm{conf}}$ is plotted against normalized total cell count $\eta_{\mathrm{tot}}$ for CT, AT50, AT100, ATagent and the distilled transparent strategy AT$^{+}$. Experimental trajectories (Exp, left) are compared with the corresponding spatial simulations (Sim, right), with points coloured by the normalized resistant-cell count $\eta_{\mathrm{res}}$.

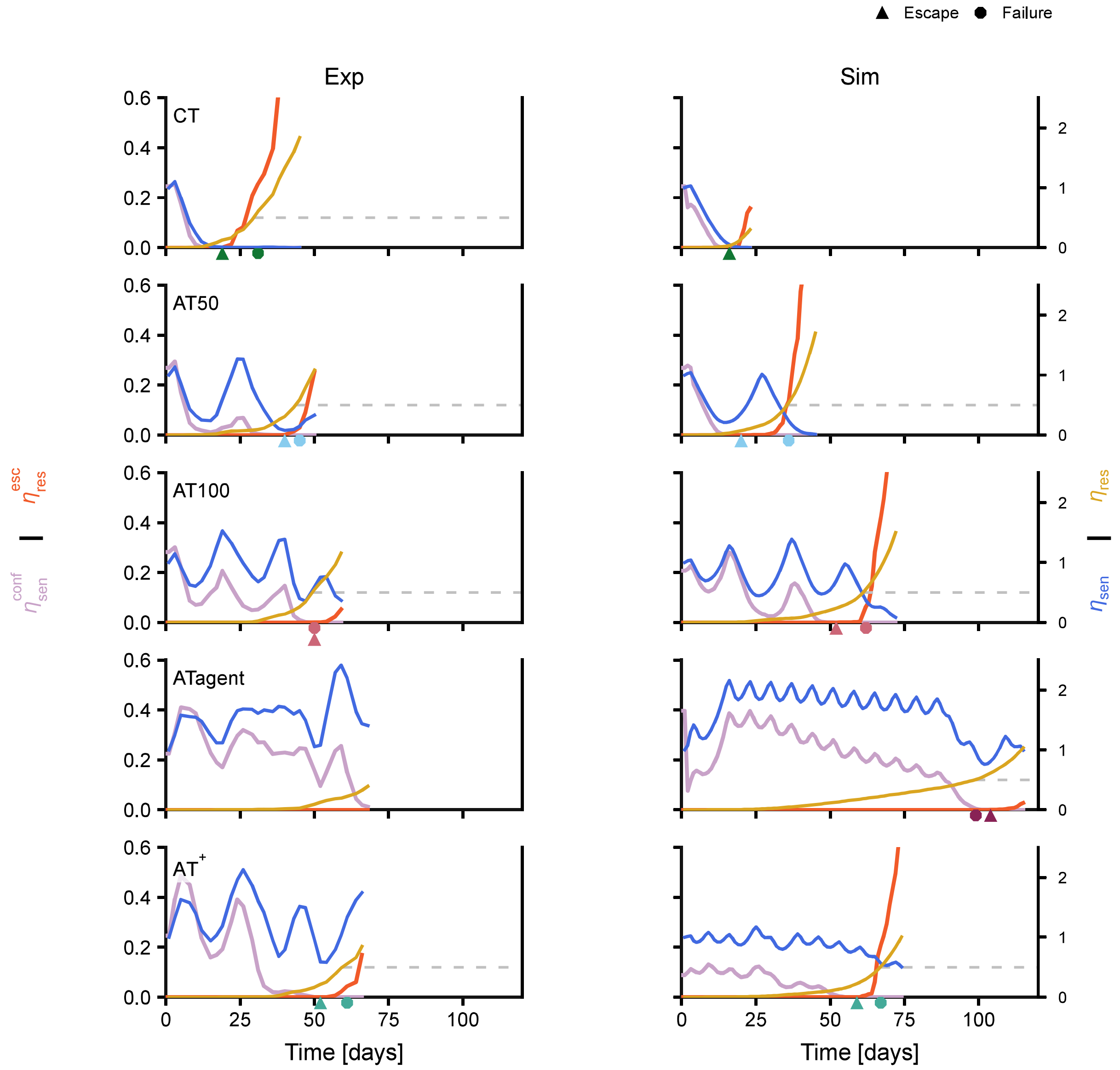

Supplementary Figure 6: **Escape-margin dynamics across treatment strategies.** Confinement and escape components of the escape margin over time for CT, AT50, AT100, ATagent, and the distilled transparent strategy AT$^+$. The confinement component ($\eta_{\rm sen}^{\rm conf}$) and escape component ($\eta_{\rm res}^{\rm esc}$) are shown as well as the normalized sensitive and resistant cell burdens ($\eta_{\rm sen}$, $\eta_{\rm res}$). Experimental trajectories (Exp, left) are compared with the corresponding simulated trajectories (Sim, right). Gray line indicates treatment failure threshold.

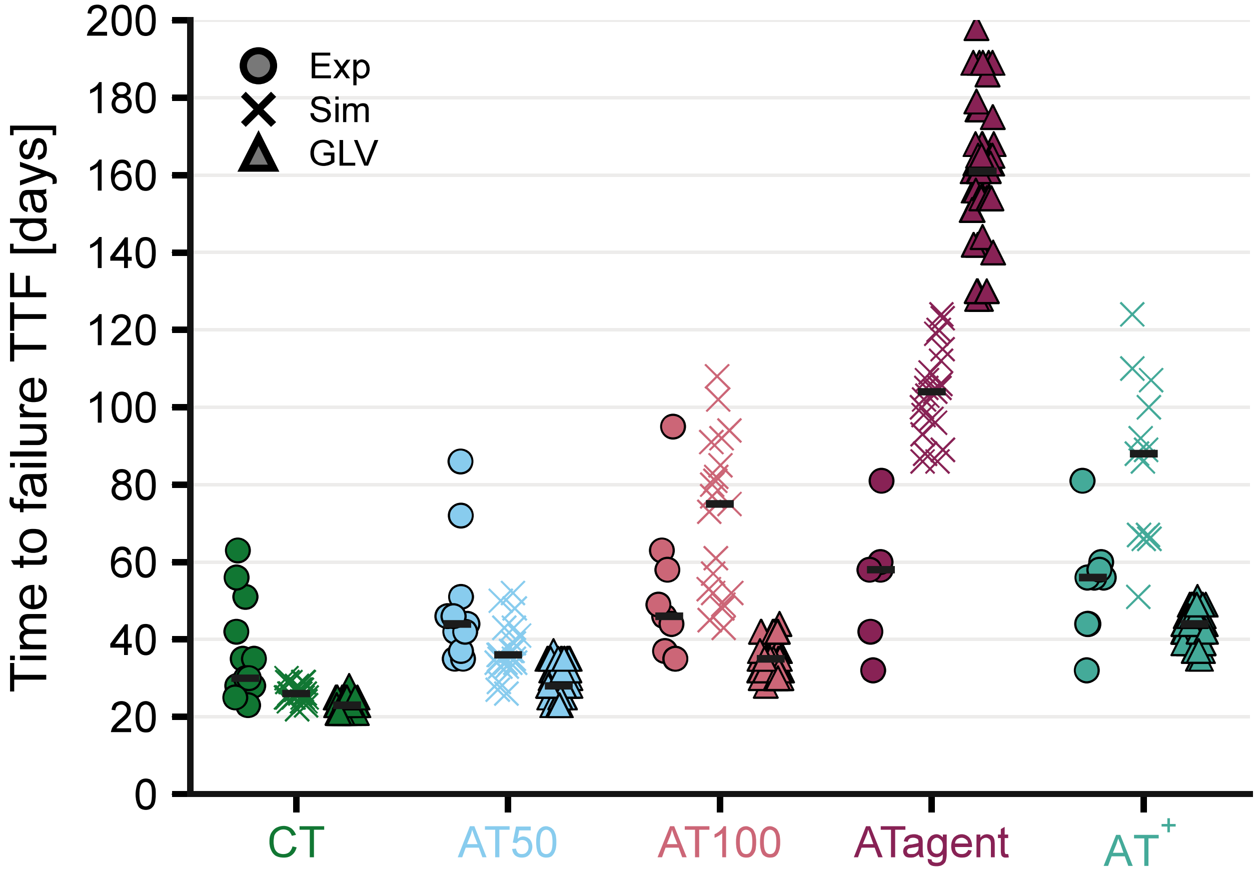

Supplementary Figure 7: **Time to failure across treatment strategies and model systems.** Time to failure (TTF) is shown for CT, AT50, AT100, ATagent and the distilled transparent strategy $AT^{+}$. Results are shown for experimental replicates (Exp, circles), spatial simulations (Sim, crosses), and the generalized Lotka–Volterra model (GLV, triangles). Individual points represent replicate trajectories.

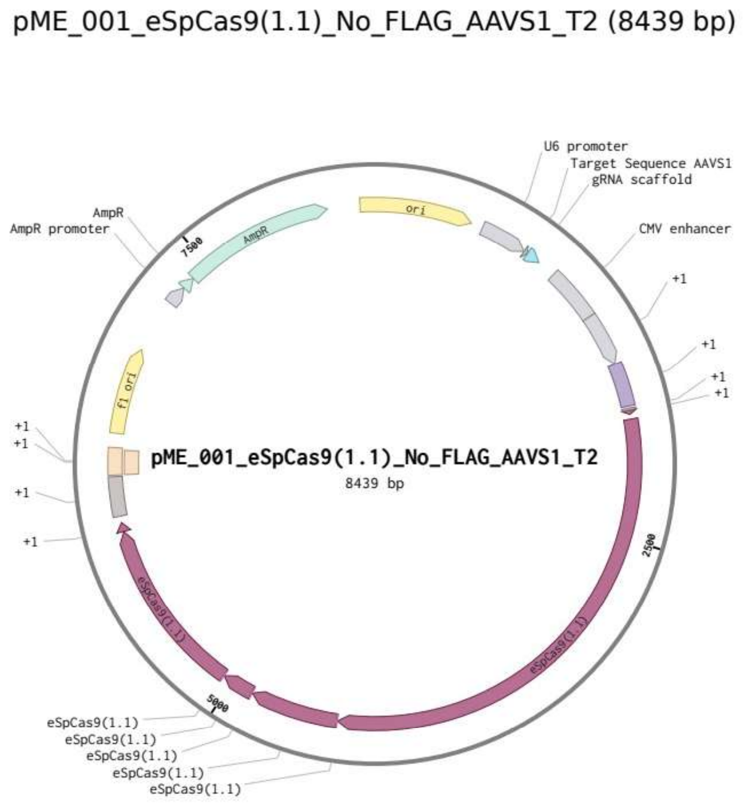

Supplementary Figure 8: **Plasmid map of the AAVS1-targeting construct pME_001.** The 8,439-bp plasmid encodes eSpCas9(1.1) and a U6-driven sgRNA targeting the AAVS1-T2 site. Selected backbone elements, including the bacterial origin of replication and ampicillin-resistance cassette, are indicated.

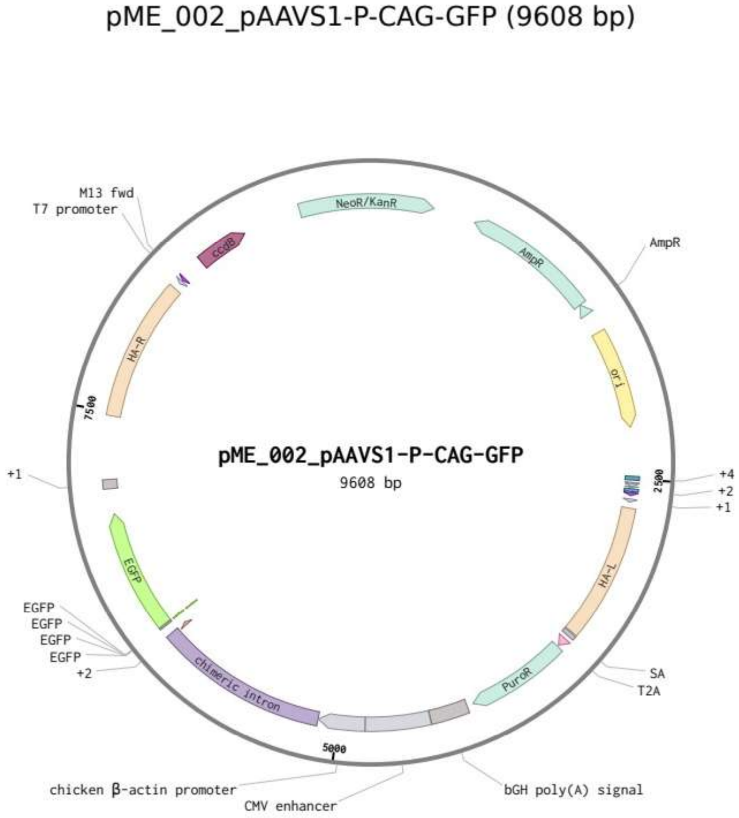

Supplementary Figure 9: **Plasmid map of the EGFP donor construct pME_002.** The 9,608-bp AAVS1 donor plasmid contains left and right AAVS1 homology arms, a CAG-driven EGFP expression cassette, and a promoterless splice-acceptor-T2A-puromycin-resistance cassette. Selected bacterial backbone elements are indicated.

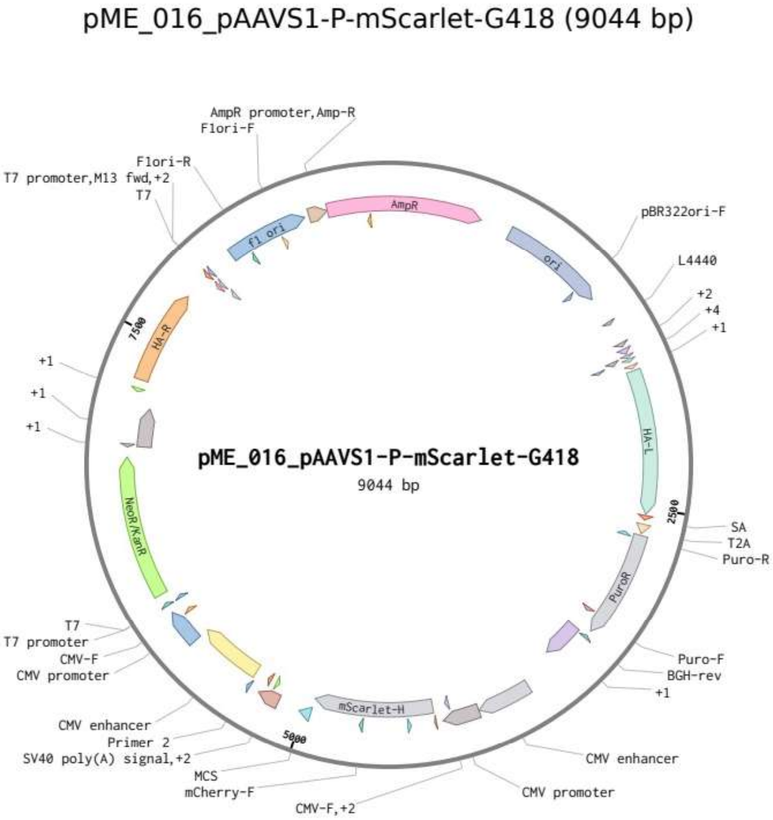

Supplementary Figure 10: **Plasmid map of the mScarlet and G418-resistance donor construct pME_016.** The 9,044-bp AAVS1 donor plasmid contains left and right AAVS1 homology arms, an mScarlet-H expression cassette, a CMV-driven neomycin-resistance cassette conferring G418 resistance, and a promoterless splice-acceptor-T2A-puromycin-resistance cassette. Selected bacterial backbone elements are indicated.

### Supplementary Tables

| Primer pair | Forward primer sequence (5'–3') | Reverse primer sequence (5'–3') | Application/product |
|---|---|---|---|
| oME_095 / oME_096 | GAGCCATCTCTCTCCTTGCC | GTTAGACCCAATATCAGGAGAC | Amplification of the pME_012-derived AAVS1 donor backbone (5,400 bp) |
| oME_161 / oME_173 | CTCCTGATATTGGGTCTAACCTGATTCTGTGGATAACC | ATCCGTCGACGTCAGGTGGCCTTACAATTTACGCGTTAAG | Amplification of the mScarlet-H cassette (1,670 bp) |
| oME_174 / oME_105 | GCCACCTGACGTCGACGG | GGCAAGGAGAGAGATGGCTCGCTGCGCGTAACCACCAC | Amplification of the NeoR cassette (1,900 bp) |
| oME_175 / oME_176 | GAACAGTACGAACGCTCCGAG | GTCATAGCCGAATAGCCTCTCC | Colony PCR screening of pME_016 |
| oME_066 / oME_067 | TGGAGGAGAATCCACCCAAAAG | TGCCCTGGCTCACAAATACCA | AAVS1 integration control for sensitive cells |
| oME_070 / oME_071 | TCGTCCGCGACCCACA | ATTGTCACTTTGCGCTGCC | AAVS1 integration control for sensitive cells |
| oME_068 / oME_264 | GGCTCCATCGTAAGCAAACC | CAGGACATAGCGTTGGCTAC | AAVS1 insertion control for resistant cells (572 bp) |
| oME_068 / oME_175 | GGCTCCATCGTAAGCAAACC | GAACAGTACGAACGCTCCGAG | Amplification across the complete NeoR sequence (2,400 bp) |
| oME_068 / oME_069 | GGCTCCATCGTAAGCAAACC | CCTTCTCCGACGGATGTCTC | AAVS1 locus control (622 bp) |
| oME_039 / oME_040 | GTGAGACTTGGAACCTGGGG | TCCTGCAGCAGTGGAATCAG | Predicted off-target locus 1 (516 bp) |
| oME_041 / oME_042 | GAGAGATGCTCTGGGCTTCC | GGAGCCACCTTGCAGAAGAT | Predicted off-target locus 2 (592 bp) |
| oME_043 / oME_044 | TCCTTAAGCCCCAAAGCTGG | GGGGATGGGAGGGTTACTCT | Predicted off-target locus 3 (700 bp) |

Table S2: **Primers used for plasmid construction and validation of the engineered cell lines.** Sequences are given in the 5'–3' direction. For primers used in Gibson assembly, sequences include the complete homologous overlaps.

| Plate | Complete exclusions | Endpoint cutoffs (final retained day) |
|---|---|---|
| 1 | A1–A6 (AT50); B1 (AT100); C1 (Pulse); D1, D2, D3, D5, D6 (NT) | B3 (AT100; d38); B4 (AT100; d47); B5 (AT100; d22) |
| 2 | A2, A4, A5 (CT); B3 (AT50); C3 (AT100); D1, D2, D3, D6 (Pulse) | A1 (CT; d33); A3 (CT; d47); A6 (CT; d36); B1 (AT50; d33); B2 (AT50; d50); C2 (AT100; d36); C4 (AT100; d40); C5 (AT100; d45); C6 (AT100; d38) |
| 3 | A1–A6 (NT); B1, B4, B5, B6 (CT); C1, C6 (AT50); D1, D2, D3, D5, D6 (AT100) | B2 (CT; d45); B3 (CT; d94); C3 (AT50; d38); C5 (AT50; d89); D4 (AT100; d47) |
| 4 | A1, A2, A3, A6 (Pulse); C3, C6 (CT) | C1 (CT; d50); C2 (CT; d50); C4 (CT; d68); D1 (AT50; d36); D2 (AT50; d50); D3 (AT50; d45); D4 (AT50; d43); D5 (AT50; d54) |
| 5 | A1, A3, A6 (AT100); B6 (Pulse); D1, D6 (CT) | A2 (AT100; d43); A5 (AT100; d59); D2 (CT; d47) |
| 6 | A1, A2, A6, C1, C2, C5, C6 (ATAgent1); B1, B6, D1–D6 (ATAgent2) | A5 (ATAgent1; d47); B2 (ATAgent2; d68); B3 (ATAgent2; d61); B4 (ATAgent2; d68); B5 (ATAgent2; d54) |
| 7 | B3, D1–D6 (ATAgent1); A1, A3, A6, C3, C6 (ATAgent2) | A2 (ATAgent2; d52); A4 (ATAgent2; d71); A5 (ATAgent2; d40); B2 (ATAgent1; d47); B5 (ATAgent1; d52); B6 (ATAgent1; d59); C2 (ATAgent2; d68); C4 (ATAgent2; d68); C5 (ATAgent2; d36) |
| 8 | A1–A6, B1, B5, B6, C1, C6, D1, D2, D6 (Dens) | B2 (Dens; d47); B4 (Dens; d57); C2 (Dens; d54); C3 (Dens; d64); C4 (Dens; d52); C5 (Dens; d38); D3 (Dens; d66); D4 (Dens; d61); D5 (Dens; d66) |

Table S3: **Manual replicate exclusions and analysis cutoffs.** Replicate identifiers are abbreviated as well (protocol); the complete identifier used in the analysis is `well_protocol-well`. “dXX” denotes the final day retained for a replicate subject to an endpoint cutoff. The table reproduces the shared filtering rules used by the analysis scripts. These exclusions apply to spatial and outcome analyses; effective-model calibration and bootstrap refitting used a separate fixed dataset, including colonies completely excluded from those analyses.
